# The chemotherapy-induced senescence-associated secretome promotes cell detachment and metastatic dissemination through metabolic reprogramming

**DOI:** 10.1101/2023.12.02.569652

**Authors:** Aidan R. Cole, Raquel Buj, Apoorva Uboveja, Evan Levasseur, Hui Wang, Katarzyna M. Kedziora, Adam Chatoff, Andrea Andress Huacachino, Mariola M. Marcinkiewicz, Amandine Amalric, Baixue Yang, Naveen Kumar Tangudu, Jeff Danielson, Amal Elwah, Sierra White, Danyang Li, Callen T. Wallace, Felicia Lazure, Esther Elishaev, Lauren Borho, Dorota E. Jazwinska, Matthew S. Laird, Huda Atiya, Benjamin G. Bitler, Denarda Dangaj, Lan G. Coffman, George Tseng, Steffi Oesterreich, Ana P. Gomes, Aditi U. Gurkar, Francisco J. Schopfer, Francesmary Modugno, Simon C. Watkins, Ioannis K. Zervantonakis, Wayne Stallaert, Nadine Hempel, Nathaniel W. Snyder, Katherine M. Aird

**Affiliations:** Department of Pharmacology & Chemical Biology and UPMC Hillman Cancer Center, University of Pittsburgh School of Medicine, Pittsburgh, PA; Department of Cell Biology, Center for Biologic Imaging, University of Pittsburgh School of Medicine, Pittsburgh, PA; Aging + Cardiovascular Discovery Center, Department of Cardiovascular Sciences, Lewis Katz School of Medicine, Temple University, Philadelphia, PA; Tsinghua University School of Medicine, Beijing, P.R. China; Department of Medicine and UPMC Hillman Cancer Center, University of Pittsburgh, Pittsburgh, PA; Department of Biostatistics, University of Pittsburgh, Pittsburgh, PA; Department of Molecular Oncology, H. Lee Moffitt Cancer Center & Research Institute, Tampa, FL; Department of Pathology, University of Pittsburgh School of Medicine, Pittsburgh, PA; Department of Obstetrics, Gynecology and Reproductive Sciences, University of Pittsburgh School of Medicine and Magee-Womens Research Institute, University of Pittsburgh, Pittsburgh, PA; Department of Bioengineering and UPMC Hillman Cancer Center, University of Pittsburgh, Pitts-burgh, PA; Division of Malignant Hematology & Medical Oncology, Department of Medicine, UPMC Hillman Cancer Center, University of Pittsburgh School of Medicine, Pittsburgh, PA; Division of Reproductive Sciences, University of Colorado Anschutz Medical Campus, Aurora, CO; Ludwig Institute for Cancer Research, University of Lausanne (UNIL), Lausanne, Switzerland; Aging Institute of UPMC and the Division of Geriatric Medicine, Department of Medicine, University of Pittsburgh School of Medicine, Pittsburgh, PA; Department of Pharmacology and Chemical Biology, University of Pittsburgh, Pittsburgh, PA, U.S.A; Pittsburgh Heart, Lung, Blood, and Vascular Medicine Institute (VMI); Pittsburgh Liver Research Center (PLRC); and Center for Immunometabolism, Pittsburgh, PA; Department of Computational and Systems Biology, UPMC Hillman Cancer Center, University of Pittsburgh, PA

**Keywords:** complex I, NDUFA5, oxidative phosphorylation, NAD, SREBP, sirtuins, cholesterol, glucose, fructose, metastasis

## Abstract

Cellular senescence, characterized by a stable cell cycle arrest, is a well-documented consequence of several widely used chemotherapeutics that has context-dependent roles in cancer. Although senescent cells are non-proliferative, they remain biologically active and secrete a complex and diverse array of factors collectively known as the se-nescence-associated secretome (SAS), which exerts pro-tumorigenic effects. Here, we aimed to mechanistically investigate how the SAS contributes to metastatic dissemination of high grade serous ovarian cancer (HGSOC) using standard-of-care cisplatin as a senescence inducer. Our findings demonstrate that the cisplatin-induced SAS enhances the dissemination of HGSOC *in vivo* without affecting cell proliferation or viability. We found that the SAS facilitates cell detachment, an effect that is mediated by a metabolic component. Using a metabolically focused CRISPR knockout screen, we identified complex I as the key driver of SAS-mediated cell detachment in bystander cells and validated that inhibition of complex I activity decreases HGSOC dissemination *in vivo*. Mechanistically, this effect was driven by SAS-mediated inhibition of an NAD^+^-SIRT-SREBP axis, leading to decreased plasma membrane cholesterol that increased cell detachment. Excitingly, we found that fructose is the key SAS component upstream of the NAD^+^-SIRT-SREBP-cholesterol axis mediating increased detachment of bystander cells, and a high fructose diet increases HGSOC dissemination *in vivo*. These findings reveal that the cisplatin-induced SAS reprograms the metabolic microenvironment in HGSOC, driving cancer cell detachment and promoting metastatic dissemination in a paracrine fashion. They also point to a previously unrecognized pro-tumorigenic effect of the SAS that may contribute to the high recurrence rate of HGSOC patients.

## INTRODUCTION

Cellular senescence is defined as a stable cell cycle arrest^1,2^. In cancer, senescence is a common cell fate induced by chemotherapy (termed therapy-induced senescence: TIS)^3–5^. TIS was initially thought to be a beneficial therapeutic response^6–8^; however, over the past decade this paradigm has shifted, and multiple papers have reported a more nuanced and complex role for TIS in cancer. Recent evidence has demonstrated that TIS paradoxically enhances tumorigenesis, metastasis, and chemoresistance in a paracrine fashion^8–14^. The pro-tumorigenic role for TIS in cancer is due in part to the unique secretory phenotype of senescent cells (termed the senescence-associated secretory phenotype: SASP). This secretome, herein referred to as the senescence associated secretome (SAS), is composed of a variety of factors and is different depending on both the senescence inducer and cell type^15–18^. The SASP is often described as a transcriptional program that alters the inflammatory phenotype of senescent cells and their tumor microenvironment (TME)^19,20^. However, SAS factors go beyond transcriptionally regulated inflammatory mediators, and there is evidence that metabolites such as bioactive lipids, fatty acids, nucleobases, amino acids, and others are SAS components^21–23^. Moreover, senescent cells reprogram their intracellular metabolism^24,25^, and a few publications have pointed to depletion of certain metabolites in the SAS^21,26^. Yet the metabolic components of the SAS remain underexplored, and it is still unknown how these metabolites influence non-senescent bystander cancer cells within the TME.

Altered metabolism is hallmark of cancer that is known to drive metastasis in several cancer types^27–29^. However, the extracellular signals leading to these intracellular metabolic changes remain underexplored. One key characteristic of metabolic reprogramming in cancer is through changes in mitochondrial metabolism, including differential electron transport chain (ETC) activity^30^. Recently, the role of complex I of the ETC has been identified as a driver of metastasis through downstream mechanisms involving NAD^+^ regeneration, ROS, and/or energy production^31–33^. Even with the recent advances in understanding senescent cell metabolism and metabolic components of the SAS, whether and how metabolic reprogramming from the SAS contributes to bystander cancer cell metabolic reprogramming and subsequent metastasis has not been fully explored.

One chemotherapy well-known to promote TIS is cisplatin^34,35^. This is of particular importance in high grade serous ovarian cancer (HGSOC), where platinum-based therapies such as cisplatin are standard-of-care^36^. While the majority of HGSOC patients initially respond to platinum-based chemotherapy, disease recurrence is common, accounting for ∼90% of HGSOC deaths^37^. The use of chemotherapy is associated with elevated metastatic dissemination of ovarian cancer^38,39^, and recurrent HGSOC patients all have disseminated disease^40^. Metastatic dissemination of HGSOC occurs most frequently through the transcoelomic route, defined as shedding of cancer cells from a solid tumor (either the primary tumor or peritoneal tumor nodules) into the peritoneal cavity^41,42^. The first step in transcoelomic dissemination is detachment of cells from the primary tumor^43–46^. While this initial dissociation step of this dissemination route is considered “passive”, it is associated with the loss of cell-cell adhesion factors^47–52^. The role of the cisplatin-induced SAS in cell-cell adhesion, detachment, and dissemination remains unknown. The use of HGSOC is an ideal model to investigate these significant questions in cancer senescence.

Here we aimed to investigate the paracrine effects of the cisplatin-induced SAS produced by HGSOC cells on metastatic dissemination. We found that the resulting cisplatin-induced SAS was sufficient to weaken adhesion and promote detachment of bystander cancer cells, enhancing metastatic dissemination *in vivo*. Strikingly, this effect was driven by a metabolic—rather than protein—component of the SAS. Using a metabolism-focused CRISPR knockout screen, we identified complex I, particularly NDUFA5, as a key mediator of this response. Mechanistically, SAS-driven complex I activity reduced membrane cholesterol by inhibiting NAD⁺-SIRT-SREBP signaling, leading to increased detachment in bystander cancer cells. Unexpectedly, elevated fructose in the SAS activated this pathway, and a high fructose diet enhanced dissemination *in vivo*. Clinically, high senescence signatures, high complex I signatures, and high circulating fructose levels correlated with more advanced HGSOC stage at diagnosis, indicating an association between this pathway and widespread metastasis. Together, these findings reveal a novel metabolic axis within the SAS that reprograms bystander cell metabolism to promote metastatic dissemination.

## RESULTS

### The cisplatin-induced senescence-associated secretome promotes HGSOC dissemination *in vivo*

Platinum-based therapies such as cisplatin are standard-of-care for HGSOC patients, although the majority of HGSOCs initially treated with platinum will recur with widespread disseminated disease^38,39^. Cisplatin induces senescence and a senescence-associated secretome (SAS)^34,35^, but the contribution of the SAS on dissemination and tumor progression in HGSOC remains unclear. Here, we aimed to determine the paracrine effects of the cisplatin-induced SAS on HGSOC bystander cells (**Fig. 1A**). Towards this goal, we induced senescence in two different HGSOC cell lines using cisplatin (**Fig. S1A**). Senescence in HGSOC cells was confirmed using multiple markers (**Fig. S1B-G**). To determine whether the SAS affects HGSOC progression *in vivo*, we co-injected naïve proliferating HGSOC cells with vehicle treated or cisplatin-induced senescent HGSOC cells (**Fig. 1B**). It is important to note that we injected the same number of proliferating cells in both groups (see Methods for more details). We observed an increase in tumor burden in mice co-injected with naïve proliferating cells + senescent cells compared to naïve proliferating cells + vehicle control cells (**Fig. 1C**). Moreover, we noted a metastatic tropism to the omentum, intestines, visceral fat, and peritoneum (**Fig. 1D**), while other sites were not markedly affected (**Fig. S2A**). The increase in tumor burden was not due to senescent cells regaining proliferative capacity *in vivo* (**Fig. S2B**). To more directly determine the contribution of the SAS to this increase in tumor burden, used a conditioned media approach. We treated KPCA.B HGSOC cells with vehicle or cisplatin to induce senes-cence, washed off the drug, and conditioned media for an additional 2 days (**Fig. S1A**). We then injected conditioned media from cisplatin-induced senescent HGSOC cells (SCM) or proliferating controls (PCM) into tumor-bearing mice (**Fig. 1E**). SCM alone was sufficient to increase tumor burden and metastatic spread mainly to the intestines, alt-hough there was a trend in similar tropism as the co-injection experiment (**Fig. 1F-G and S2C**). There was no marked difference in omental weights or Ki67 staining in either model (**Fig. 1H-I**), demonstrating that the observed increase in tumor nodules was likely not due to increased proliferation of cancer cells. Consistently, SCM did not increase bystander HGSOC cell proliferation *in vitro* (**Fig. S2D**). Additionally, we did not observe differences in TUNEL staining in the tumors (**Fig. 1J**), and SCM did not affect anoikis *in vitro* (**Fig. S2E**), demonstrating that the SAS does not affect the viability of HGSOC cells in suspen-sion. Finally, we used TCGA data from HGSOC tumors and found that a high senescence signature^53^ is associated with advanced stages (a proxy for disseminated disease), with nearly 100% of patients with a high senescence signature having stage IIIC or IV disease compared with 77% of patients with a low senescence signature (**Fig. 1K and Table S1**). This indicates a pro-tumorigenic role for senescence in HGSOC patients that is associ-ated with higher stage of disease. Together, these data demonstrate that the SAS from cisplatin-induced senescent HGSOC cells is sufficient to increase HGSOC dissemination *in vivo* and this is not due to either increased proliferation or decreased cell death.

**Figure 1.**
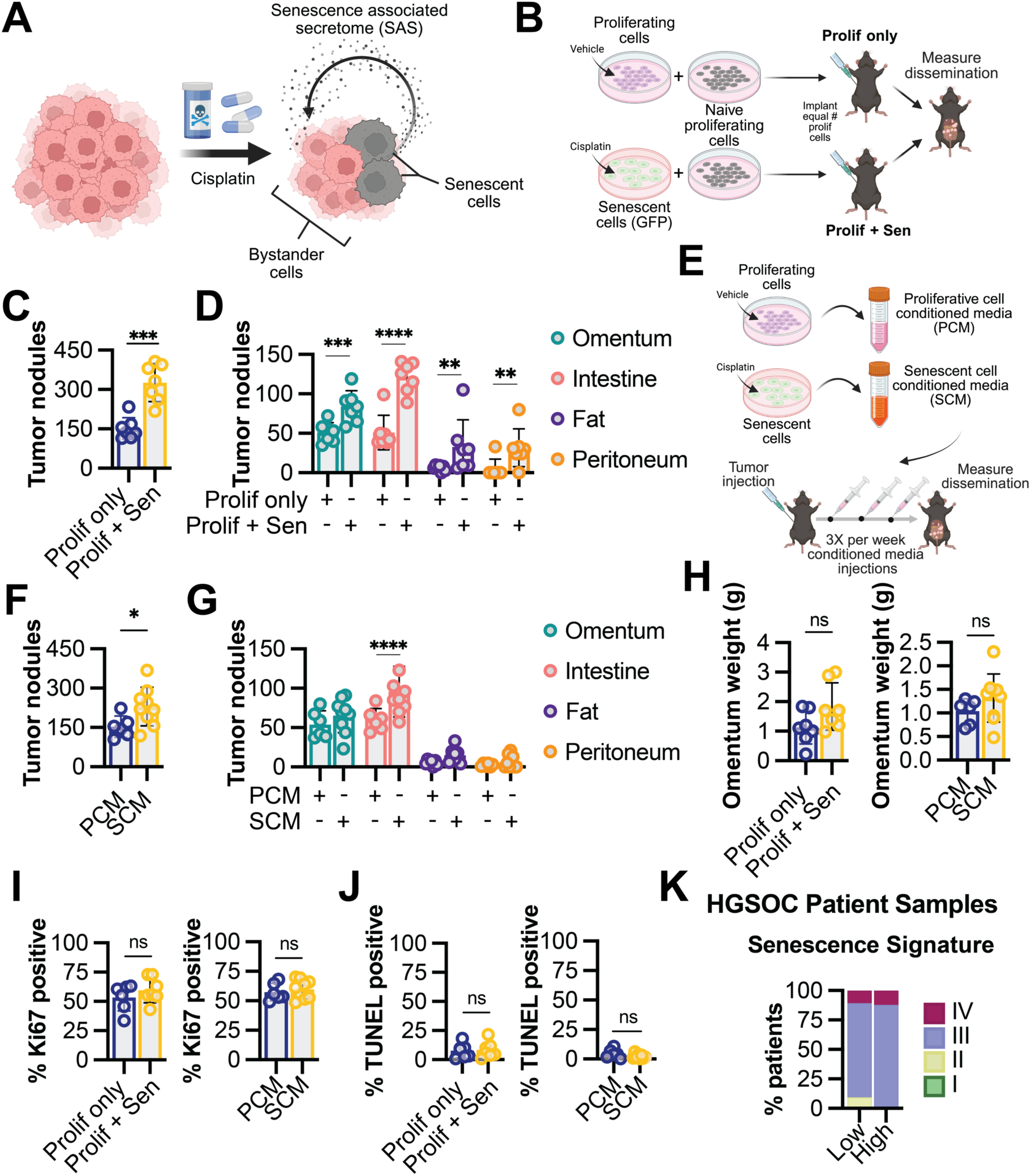
The cisplatin-induced senescence-associated secretome (SAS) promotes dissemina-tion of high grade serous ovarian cancer (HGSOC) *in vivo*. (**A**) Schematic of the post-cisplatin tumor microenvironment where senescent cells occur and can act on bystander cells in a paracrine manner through the senescence associated secretome (SAS). **(B)** Schematic of *in vivo* mouse exper-iment where control or cisplatin-induced senescent cells were co-injected with naïve proliferating cells. KPCA.B HGSOC cells were treated with vehicle or induced to senesce using cisplatin (**see Fig. S1 for timeline and senescence markers**). Cisplatin-induced senescent cells were also labeled with GFP. Vehicle treated or cisplatin-induced senescent cells were then co-injected intraperitoneally with naïve proliferating KPCA.B HGSOC. The same number of proliferating cells were injected into mice in each group, and the senescent cells were injected with naïve proliferating cells at a 1:1 ratio. Prolif only = vehicle treated cells co-injected with naïve proliferating cells. Prolif + Sen = cisplatin-induced senes-cent cells co-injected with naïve proliferating cells. **(C)** Endpoint total intraperitoneal tumor nodules from the experiment detailed in **(B)**. 6-7 mice/group. **(D)** Endpoint tumor nodule counts in the indicated metastatic sites from the experiment detailed in **(B)**. 6-7 mice/group. **(E)** Schematic of conditioned media injection *in vivo* model. KPCA.B HGSOC cells were treated with vehicle or induced to senesce using cisplatin, and conditioned media was harvested (see Methods for more details). PCM = prolifer-ative conditioned media, which is conditioned media harvested from vehicle treated cells. SCM = se-nescent conditioned media, which is conditioned media harvested from cisplatin-induced senescent cells. KPCA.B HGSOC cells were injected intraperitoneally into mice, and upon tumor establishment, conditioned media was injected 3X per week. **(F)** Endpoint total intraperitoneal tumor nodules from the experiment detailed in **(E)**. 6-9 mice/group. **(G)** Endpoint tumor nodule counts in the indicated meta-static sites from the experiment detailed in **(E)**. 6-9 mice/group. **(H)** Omental weights from *in vivo* ex-periments detailed in (B-left) and (E-right). **(I)** Quantification of Ki67 immunohistology staining of omental tumors detailed in (B-left) and (E-right). **(J)** Quantification of TUNEL immunohistology stain-ing of omental tumors detailed in (B-left) and (E-right). **(K)** HGSOC patients from TCGA were stratified based on the SenPy senescence signature^53^ (see **Table S1** for gene list). Shown is the stage of pa-tients in the lower and upper quartile. All graphs show mean ± SD; Student’s t-test; ns = not significant, *p>0.05, *p<0.05, *p<0.01, ***p<0.001, ****p<0.0001.

### The cisplatin-induced senescence-associated secretome promotes detachment of bystander HGSOC cells via NDUFA5

We found that the SAS increases the number of tumor nodules without corresponding changes in cancer cell proliferation or death (**Fig. 1 and S2**), suggesting that the SAS influences metastatic dissemination. During transcoelomic dissemination within the peritoneal cavity, HGSOC cells must detach from the solid tumor by loosening their adhesion to other cells and extracellular matrix (ECM)^41,54^. Thus, we next investigated how the cis-platin-induced SAS from HGSOC cells influences detachment and adhesion of bystander HGSOC cells. To measure detachment, we performed timelapse imaging of spheroids cultured in PCM or SCM. Spheroids cultured in round bottom ULA wells were of similar size regardless of whether they were exposed to PCM or SCM, and we did not observe differential compaction between these conditions (**Fig. S3A**), demonstrating that HGSOC cells in SCM remain able to form spheroids of the same size as those in PCM under gravitational constraint. Thus, for live cell imaging experiments, we controlled for cell number and spheroid size by forming spheroids in round bottom ultra-low attachment (ULA) wells before transferring them to flat bottom ULA wells (**Fig. 2A**). After transferring spheroids to flat bottom ULA wells, we observed an increase in the total number of live cell detachment events when bystander cells were cultured in SCM compared to PCM (**Fig. 2B-C and Video S1**), indicating that cells were more loosely adherent. Staining with ethidium homodimer confirmed the detached cells were viable (**Fig. 2B**). Interestingly, heat inactivating the SCM to denature proteins also drove detachment of live cells to the same level as regular SCM (**Fig. 2C**), demonstrating that the observed increase in detachment was likely not due to protein activity, including ECM degraders such as matrix metalloproteases (MMPs), well-known components of the SAS ^55–57^. As a complementary approach, we assessed detachment using a controlled trypsinization assay (**Fig. 2D**), similar to a previous publication^58^. Consistent with our live cell imaging results in spheroids (**Fig. 2C**), we found that bystander cells cultured in SCM had increased detachment compared to those cultured in PCM, and the increase in detachment was also retained in heat inactivated SCM (**Fig. 2E**). While most prior work has focused on the protein content of the SAS, we and others have shown that the SAS can also have differential metabolite components^21–23,26^. Therefore, we aimed to determine whether the observed phenotype is driven by a small molecule or metabolite by filtering the SCM through a filter to remove most proteins (>3kD), leaving small molecules such as metabolites. Interestingly, by-stander HGSOC cells cultured in filtered SCM also displayed increased detachment to the same level as regular SCM (**Fig. 2E**). Together, these data provide evidence that a metabolic component of the cisplatin-induced SAS may drive the observed increased de-tachment of bystander HGSOC cells.

**Figure 2.**
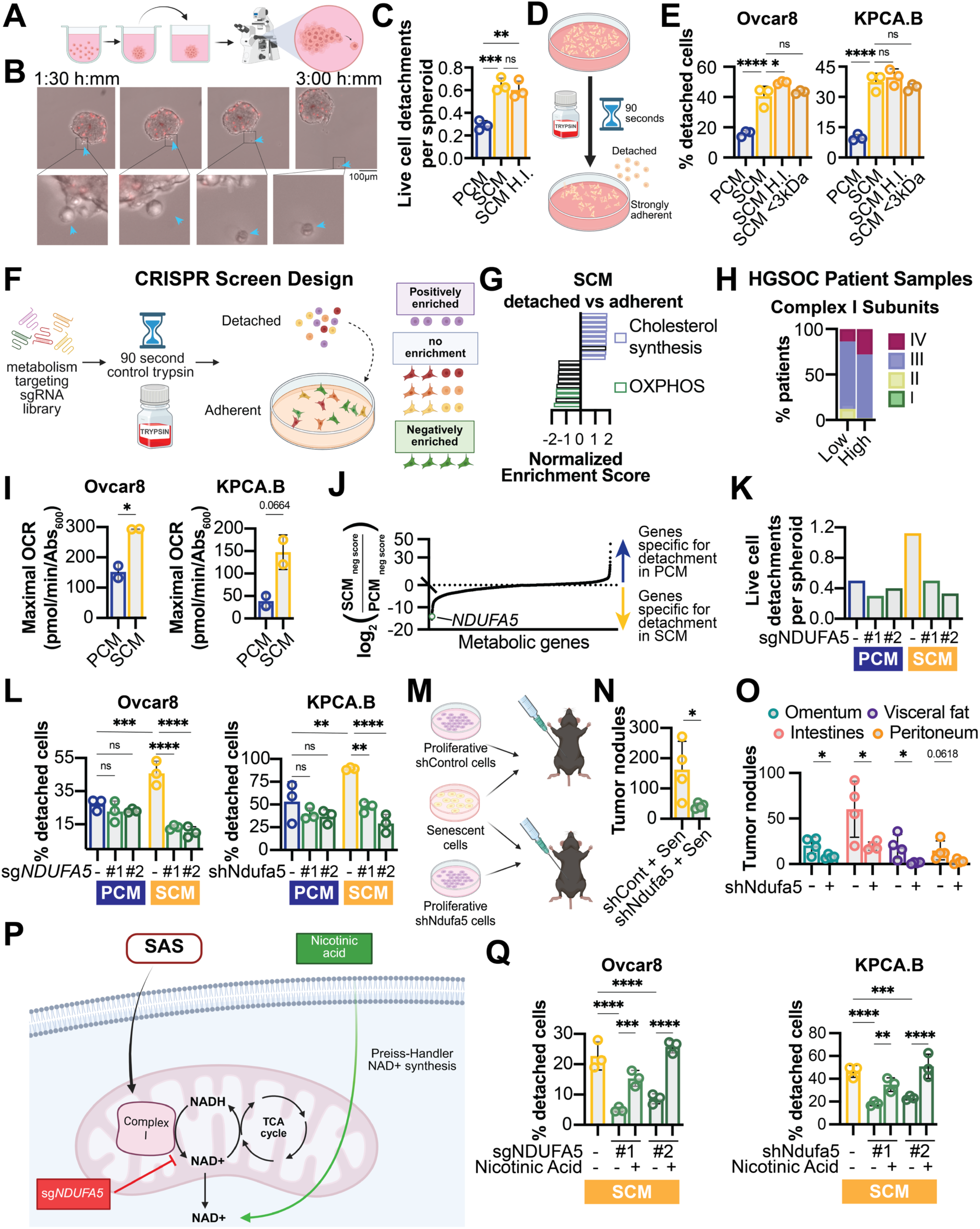
The cisplatin-induced senescence-associated secretome promotes cell detachment via NDUFA5. **(A)** Schematic of assay to assess live cell detachment from spheroids. Ovcar8 HGSOC cells were cultured in round bottom ultra-low attachment (ULA) plates to form spheroids. These spheroids were subsequently moved to flat bottom ULA plates and cultured for 48h in either vehicle treated proliferative conditioned media (PCM) or cisplatin-induced senescent conditioned media (SCM). Live cell imaging on ethidium homodimer-stained cells was performed for 24h to capture detachment events. **(B)** Representative image from live cell imaging experiment detailed in **(A)**. Blue arrows point to a live (ethidium homodimer negative) cell detaching from the main spheroid. **(C)** Quantification of live cell imaging experiment detailed in **(A)**. In addition to PCM and SCM, spheroids were also cultured in SCM that was heat inactivated (SCM H.I.) to denature proteins. n = 3 wells with 5-8 independent spheroids; one-way ANOVA. **(D)** Schematic of controlled trypsin experiments detailed in (E, F-G, J, L, Q). **(E)** The indicated bystander cells were cultured in PCM, SCM, or SCM that was heat inactivated (SCM H.I.) or filtered using a 3kD cut-off column (SCM <3kd). After 72h, the controlled trypsin experiment (detailed in D) was performed. Shown are % detached cells. One-way ANOVA. **(F)** Schematic of CRISPR KO screen to identify mediators of detachment in bystander cells cultured in PCM vs. SCM. Ovcar8 HGSOC cells were transduced with a metabolically focused gRNA library and subsequently cultured in PCM or SCM. After 72h, the controlled trypsin experiment was performed, and both adherent and detached cells were harvested. **(G)** Gene set enrichment analysis (GSEA) was performed on by-stander cells cultured in SCM (adherent vs. attached). NES = normalized enrichment score. A negative NES indicates genes whose knockout decreases detachment in SCM, whereas a positive NES indicates genes whose knockout increases detachment in SCM. **(H)** Clinical stages of HGSOC patients from TCGA stratified by divided by low-quartile vs. high-quartile expression of complex I subunits (see **Table S1** for gene list). **(I)** The indicated bystander cells were cultured in PCM or SCM for 72h, and maximal respiration was quantified using a Seahorse. Student’s t-test. **(J)** Genes from CRISPR screen in (F) ranked by ratio of negative score in SCM vs. PCM. **(K)** Ovcar8 HGSOC cells with NDUFA5 knockout (sgNDUFA5 #1 and #2) were cultured as detailed in the schematic in (A). Shown are live cell detachment events per spheroid. n=12 independent spheroids. **(L)** The indicated bystander HGSOC cells with knockout of NDUFA5 (sgNDUFA5 #1 and #2) or knockdown of Ndufa5 (shNdufa5 #1 and #2) were cultured in PCM or SCM. After 72h, the controlled trypsin experiment was performed. Shown are % detached cells. One-way ANOVA. **(M)** Schematic of senescent cells with proliferative shNdufa5 cell co-injection *in vivo* model. KPCA.B HGSOC cells were treated with vehicle or induced to senesce using cisplatin. Vehicle treated or cisplatin-induced senescent cells were then co-injected intraperitoneally with naïve proliferating shNdufa5 KPCA.B HGSOC cells. The same number of proliferating cells were injected into mice in each group, and the senescent cells were injected with naïve proliferating cells at a 1:1 ratio. **(N)** Endpoint total intraperitoneal tumor nodules from the experiment detailed in **(M)**. shCont + Sen = proliferating shControl cells co-injected with senescent cells. shNdufa5 + Sen = proliferating shNdufa5 cells co-injected with senescent cells. Student’s t-test. **(O)** Endpoint tumor nodule counts in the indicated metastatic sites from the experiment detailed in **(M)**. Graph shows mean ± SD; Student’s t-test. **(P)** Schematic of the Preiss-Handler pathway of NAD+ synthesis from nicotinic acid as an alternative NAD+ source from complex I mediated NADH oxidation. **(Q)** The indicated by-stander HGSOC cells with knockout of NDUFA5 (sgNDUFA5 #1 and #2) or knockdown of Ndufa5 (shNdufa5 #1 and #2) were cultured in SCM with and without supplementation with nicotinic acid. After 72h, the controlled trypsin experiment was performed. Shown are % detached cells. One-way ANOVA. All graphs show mean ± SD. Panels C, E, I, L, Q are representative data from 3 independent experiments (n=2-3). ns = not significant, *p>0.05, *p<0.05, *p<0.01, ***p<0.001, ****p<0.0001.

Given the observed changes in detachment in heat inactivated and filtered SCM (**Fig. 2E**), we reasoned that there is a metabolic component of the SAS that likely has paracrine metabolic effects on bystander cells to decrease their adhesion. To investigate this mechanism, we conducted a CRISPR KO screen using a metabolically focused gRNA library^59^. We adopted the controlled trypsinization approach to separate the attached and detached cells from SCM and PCM cultured groups. The live cell detachment from spheroids was not optimal for this purpose because these events only led to a single cell detachment (**Fig. 2A-C**). Given the consistency in results between live cell imaging and the controlled trypsinization experiments (**Fig. 2A-E**), we used this assay for our screen. We aimed to identify genes that were negatively enriched in detached bystander cells cultured in SCM but not PCM, indicating these genes are essential for the loosely adhesive phenotype driven by the SAS (**Fig. 2F**). We found that oxidative phosphorylation genes were negatively enriched in the detached population of SCM cultured bystander cells compared to adherent cells (**Fig. 2G and Table S2 and S3**). Conversely, cholesterol biosynthesis genes were positively enriched in the detached population of SCM cultured bystander cells compared to adherent cells (**Fig. 2G and Table S2 and S3**), demonstrating that knockout of these genes drives detachment in SCM conditions. This is in stark contrast with the gene pathways identified in PCM cultured condition (**Table S3**), indicating a unique metabolic requirement for detachment driven by the cisplatin-induced SAS. Interestingly, the negative enrichment oxidative phosphorylation signature was mainly driven by complex I subunits (**Table S4**). We confirmed that complex I genes were negatively enriched specifically in detached bystander cells cultured in SCM but not in PCM (**Fig. S3B**). To show the clinical relevance of our findings, we assessed HGSOC stage based on expression of complex I or respiration genes. Indeed, high expression of complex I or respiration genes corresponds to increased HGSOC stage (**Fig. 2H, S3C, and Table S1**). This is consistent a recent report showing that ETC genes are enriched after patient-derived cells are cultured in ascites collected after chemotherapy^60^. Additionally, analysis of metastatic recurrence in a post-chemotherapy *in vivo* model identified high Nduf4l2— a complex I subunit—in malignant cells^61^. Together with our data, this indicates a key role for complex I in detachment and dissemination after chemotherapy.

Next, we aimed to validate our CRISPR screen results. Towards this goal, we used the complex I inhibitor rotenone and confirmed the necessity of functional complex I for the increased detachment phenotype of bystander cells cultured in SCM (**Fig. S3D**). Moreover, we observed an increase in both basal and maximal respiration in bystander cells cultured in SCM compared to PCM (**Fig. 2I, Fig. S3E**), indicated a cisplatin-induced SAS-mediated increase in respiration in bystander HGSOC cells. We note maximal OCR was more robustly increased in Ovcar8 cells whereas basal OCR was more robustly increased in KPCA.B cells. For further mechanistic experiments, we chose the complex I subunit NDUFA5, which was one of the top differentially negatively enriched genes in the CRISPR screen (**Fig. 2J and Fig. S3F**) and was required for the increase in SCM-mediated respiration in bystander cells (**Fig. S3G-I**). We also validated the requirement of NDUFA5 knockdown for decreased detachment in bystander HGSOC cells cultured in SCM in both HGSOC models (**Fig. 2K-L**). Notably, we were unable to grow clones with total knockout of NDUFA5 (**Fig. S3G**). Finally, we found that knockdown of Ndufa5 abrogated the increase in dissemination in the senescent cell co-injection model *in vivo* (**Fig. 2M-N**). The metastatic site tropism was also decreased by knockdown of Ndufa5 (**Fig. 2O**). Together, these data demonstrate that the cisplatin-induced SAS regulates HGSOC cell detachment and dissemination in a paracrine manner via NDUFA5.

Finally, we aimed to determine how complex I is regulating the increased detachment phenotype. Inhibition of complex I is known to produce reactive oxygen species (ROS)^62^, and ROS can induce expression of adhesion molecules^63,64^. We did not observe an increase in ROS upon NDUFA5 knockdown in bystander cells cultured in SCM, and scavenging ROS in these cells using N-acetyl-cysteine (NAC) did not decreased detachment (**Fig. S3J-K**), indicating that the SAS regulates cell adhesion via complex I activity independently of ROS generation. Complex I is also critical for oxidation of NADH to NAD^+65^ (**Fig. 2P**). Therefore, we aimed to determine whether replenishing NAD^+^ using nicotinic acid supplementation (**Fig. 2P**) would reverse the NDUFA5 knockdown-mediated decrease in detachment of bystander cells in the context of the SAS. Indeed, nicotinic acid supplementation of NDUFA5 knockdown bystander cells cultured in SCM restored the SAS-mediated increase in detachment (**Fig. 2Q**). These data point to a critical role for complex I-mediated NAD+ generation in the regulation of cell attachment in the context of the SAS.

### Bystander cell detachment is increased through inhibition of SREBP1-mediated plasma membrane cholesterol downstream of NDUFA5

We found that the complex I subunit NDUFA5 drives the cisplatin-induced SAS-mediated increased cell detachment in bystander HGSOC cells (**Fig. 2**). To understand how NDUFA5 regulates cell detachment, we performed RNA-Seq on human and murine HGSOC cells with NDUFA5 knockdown (via sgRNA or shRNA, respectively), cultured in either SCM or PCM. Pathway analysis of genes consistently upregulated by NDUFA5 knockdown in SCM, but not in PCM (**Fig. 3A and Table S5**), revealed enrichment of cholesterol-related gene signatures (**Table S6**). More specifically, *de novo* cholesterol synthesis genes were downregulated in bystander cells cultured in SCM, which was rescued by NDUFA5 knockdown (**Fig. S4A**). Consistent with the idea that cholesterol regulates detachment in bystander cells cultured in SCM, our CRISPR KO screen also identified multiple *de novo* cholesterol synthesis enzymes to be positively enriched in the detached population (**Fig. 2G and S4B**). Together, these data indicate that the SCM-driven loss of detachment in bystander cells is due to transcriptionally downregulated *de novo* cholesterol synthesis downstream of NDUFA5. Cholesterol is an important component of the plasma membrane and has been previously shown to help stabilize integrins, thereby increasing cell adhesion ^66–68^. Thus, we aimed to assess plasma membrane cholesterol content changes downstream of NDUFA5 using Filipin III, a fluorescent cholesterol probe, and total internal reflection fluorescence (TIRF) imaging. We confirmed a decrease in plasma membrane cholesterol of bystander cells cultured in SCM, which was rescued by knockdown of NDUFA5 (**Fig. 3B**). Supplementation with nicotinic acid increased the plasma membrane cholesterol content of NDUFA5 knockdown cells (**Fig. 3B**), demonstrating this effect is through NAD^+^. This was not simply due to differences in cholesterol trafficking to the plasma membrane as total cholesterol was also similarly altered (**Fig. S4C-D**). To determine whether these changes in cholesterol play a functional role in cell detachment, we depleted cholesterol with either methyl-β-cyclodextrin (MβCD), which specifically depletes plasma membrane cholesterol^69^, or statins to more generally deplete intracellular cholesterol. Both MβCD and statins abrogated the decreased detachment in NDUFA5 knockdown bystander cells cultured in SCM (**Fig. 3C and S4E-F**). Conversely, supplementation of cholesterol decreased detachment in bystander cells cultured in SCM but had no effect on NDUFA5 knockdown bystander cells cultured in SCM (**Fig. 3D**). These data demonstrate that the cisplatin-induced SAS drives differential cholesterol content in cells via NDUFA5-mediated NAD^+^, which is critical for the detachment phenotype in bystander cells.

**Figure 3.**
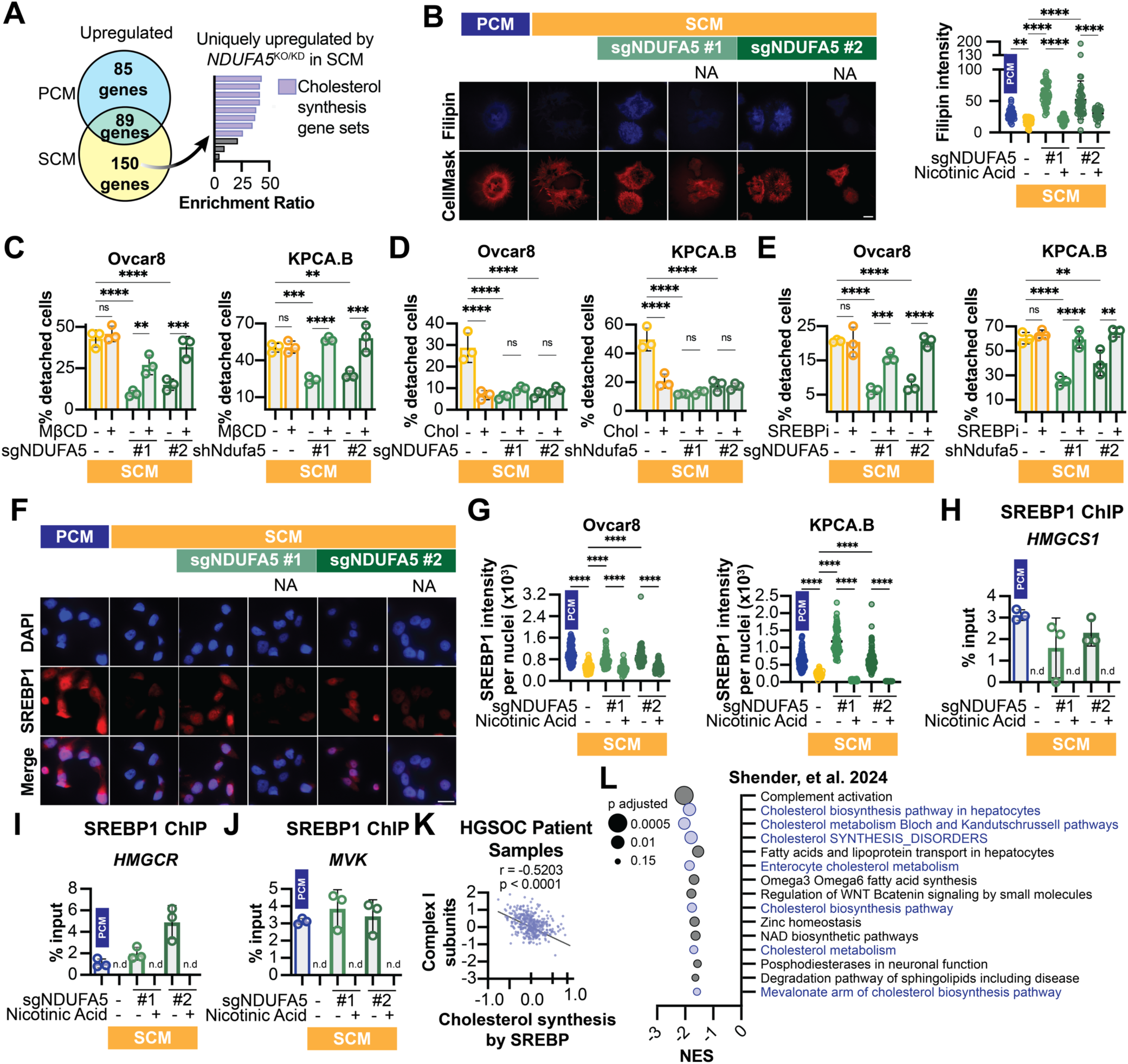
SAS-induced complex I activity via NDUFA5 increases cell detachment via decreased SREBP1-driven cholesterol synthesis. **(A)** Ovcar8 or KPCA.B bystander cells with either knockout or knockdown of NDUFA5, respectively, were cultured in PCM or SCM for 72h, and RNA was extracted. RNA-Seq was performed. Left: Venn-diagram of genes upregulated by both knockout of NDUFA5 (sgNDUFA5 #1 and #2) or knockdown of Ndufa5 (shNdufa5 #1 and #2) bystander cells cultured in PCM or SCM. Right: over-representation analysis of the genes uniquely upregulated in bystander cells cultured in SCM, but not in PCM. **(B)** The indicated bystander Ovcar8 cells with knockout of NDUFA5 (sgNDUFA5 #1 and #2) were cultured in PCM or SCM with or without nicotinic acid (NA). After 72h, plasma membrane cholesterol was assess using filipin III fluorescent staining under TIRF imaging. Left: Representative images. Scale bar = 25 µm. 100X magnification. Right: Quantification. **(C)** The indicated HGSOC bystander cells with knockout of NDUFA5 (sgNDUFA5 #1 and #2) or knockdown of Ndufa5 (shNdufa5 #1 and #2) were cultured in SCM for 72h before being treated with 10mM methyl-β-cyclodextrin (MβCD) for 1h to deplete plasma membrane cholesterol, and the controlled trypsin experiment was performed. Shown are % detached cells. **(D)** HGSOC bystander cells with knockout of NDUFA5 (sgNDUFA5 #1 and #2) or knockdown of Ndufa5 (shNdufa5 #1 and #2) were cultured in SCM for 72h before being treated with 50µM cholesterol for 2h, and the controlled trypsin experiment was performed. Shown are % detached cells. **(E)** HGSOC bystander cells with knockout of NDUFA5 (sgNDUFA5 #1 and #2) or knockdown of Ndufa5 (shNdufa5 #1 and #2) were cultured in SCM with 1µM of the SREBP inhibitor (SREBPi) fatostatin. After 72h, the controlled trypsin experiment was performed. Shown are % detached cells. **(F-G)** HGSOC bystander cells with knockout of NDUFA5 (sgNDUFA5 #1 and #2) or knockdown of Ndufa5 (shNdufa5 #1 and #2) were cultured in PCM or SCM with or without nicotinic acid supplementation. After 72h, cells were stained with anti-SREBP1 antibody and DAPI and imaged. **(F)** Representative images of SREBP1 immunofluorescent staining in Ovcar8 cells. Scale bar = 25 µm. 100X magnification. 40X magnification. **(G)** Quantification of the average SREBP1 immunofluorescent staining intensity per cell nuclei. **(H-J)** Ovcar8 bystander cells with knockout of NDUFA5 (sgNDUFA5 #1 and #2) were cultured in PCM or SCM with or without nicotinic acid. After 72h, cells were harvested for chromatin immunoprecipitation (ChIP) using an anti-SREBP1 antibody following by qPCR. Shown is occupancy of SREBP1 in the promoter of *de novo* cholesterol synthesis genes *HMGCS1* (H), *HMGCR* (I), and *MVK* (J). n.d. = not detected. **(K)** Negative correlation between the expression of genes related with complex I and cholesterol synthesis genes in TCGA HGSOC patient tumors (see **Table S1** for gene lists). Pearson’s correlation. **(L)** GSEA analysis of primary ovarian cancer cells isolated from individual patients cultured with autologous ascites fluid collected pre-vs. post-chemotherapy treatment. Wikipathways are shown. Panels D and G are representative data from 2 independent experiments in each cell line (D: n=3; G: n>100 cells). Panels B, C, E, H, I, J are representative data from 3 independent experiments (n=3). All graphs show mean ± SD; one-way ANOVA. ns = not significant, *p>0.05, *p<0.05, *p<0.01, ***p<0.001, ****p<0.0001.

We found that the changes in *de novo* cholesterol were transcriptionally driven down-stream of NDUFA5 (**Fig. 3A and S4A**), and decreased cholesterol drives the increased detachment phenotype of bystander cells cultured in cisplatin-induced SAS (**Fig. 3C-D**). Transcription of *de novo* cholesterol synthesis genes is well-known to be coordinated via the transcription factor SREBP1^70^. Next, we sought to determine whether SREBP1 contributes to the downregulation of *de novo* cholesterol synthesis genes and the increase in cell detachment downstream of NDUFA5 in SCM-cultured bystander cells. Inhibition of SREBP increased detachment specifically in NDUFA5 knockdown bystander cells cultured in SCM (**Fig. 3E**), demonstrating the requirement of SREBP1 in this phenotype. These data are also consistent with results demonstrating that cholesterol depletion increases detachment in NDUFA5 knockdown bystander cells cultured in SCM (**Fig. 3C and S4E-F**). SREBP1 translocates to the nucleus, where it binds to target genes to regulate their transcription^70^. We observed a decrease in both nuclear SREBP1 and SREBP1 occupancy on multiple *de novo* cholesterol synthesis gene promoters in bystander cells cultured in SCM that was rescued by NDUFA5 knockdown (**Fig. 3F-J and Fig. S4G**). These data are consistent with the observed decrease in *de novo* cholesterol synthesis gene expression (**Fig. 3A and S4A**). Interestingly, nicotinic acid supplementation abrogated the NDUFA5 knockdown-mediated decrease in nuclear SREBP1 and occupancy of SREBP1 at *de novo* cholesterol genes (**Fig. 3F-J**). This reinforces a role for NAD^+^ downstream of complex I activity in regulating SREBP activity. In HGSOC patient samples from TCGA, we found that expression of complex I or aerobic respiration genes inversely correlates with cholesterol synthesis genes regulated by the master transcription factor SREBP (**Fig. 3K, Fig. S4H, and Table S1**). Finally, we used a dataset in which primary ovarian cancer cells were isolated from individual patients and cultured with autologous ascites fluid collected before or after platinum plus taxane chemotherapy^60^. Interestingly, gene sets related to cholesterol synthesis were consistently downregulated in cells cultured in ascites post-chemotherapy compared to pre-chemotherapy (**Fig. 3L**). Together, these data demonstrate that downregulation of SREBP1 nuclear localization and activity by SCM decreases *de novo* cholesterol synthesis gene transcription and adhesion downstream of NDUFA5 (**Fig. S4I**).

### NDUFA5 suppresses SREBP1 via increased NAD^+^-dependent SIRT activity, leading to decreased cholesterol and detachment

We found that SCM decreases SREBP1 activity via the complex I subunit NDUFA5, leading to increased detachment in bystander cells (**Fig. 3**). Additionally, NAD^+^ pool replenishment with nicotinic acid reversed the decreased detachment caused by NDUFA5 knockdown and restored SREBP1 nuclear localization and promoter occupancy at cholesterol synthesis genes (**Fig 3F-J**), altogether demonstrating a pivotal role for NAD^+^ in this process. Replenishing NAD^+^ to SREBP-inhibited NDUFA5 knockdown bystander cells cultured in SCM did not change detachment (**Fig. 4A**), indicating that NAD^+^ is indeed upstream of SREBP1 activity. Previous reports demonstrate that SREBPs can be deactivated by sirtuins^71–73^, a family of NAD^+^-dependent deacetylases^74^. Thus, we reasoned that SIRT activity driven by NAD^+^ may increase detachment downstream of NDUFA5 by inhibiting SREBP1-mediated cholesterol synthesis. Indeed, inhibition of SIRTs prevented detachment in SCM cultured bystander cells (**Fig. 4B**). This was not due to a parallel pathway as inhibition of SIRTs did not affect detachment in NDUFA5 knockdown cells cultured in SCM (**Fig. 4B**). Addition of nicotinic acid to NDUFA5 knockdown cells cultured in SCM increased detachment, which was abrogated by inhibition of SIRTs (**Fig. 4B**). Together, these data unveil a pathway where NDUFA5 acts upstream of SIRTs via NAD^+^. To confirm that SREBP1 activity and SIRT activity are acting in the same pathway, we assessed detachment of bystander cells in response to both SIRT and SREBP inhibition. SIRT inhibition alone decreased detachment of bystander cells cultured in SCM, but not when combined with the SREBP inhibitor (**Fig. 4C**). Furthermore, we found that inhibition of SIRTs increased nuclear SREBP1 in bystander cells cultured in SCM (**Fig. 4D-E**) and increased SREBP1 occupancy at *de novo* cholesterol gene promoters (**Fig. 4F**). Finally, given the role of cholesterol in our pathway downstream of NDUFA5 and SREBP1 (**Fig. 3**), we assessed cholesterol content in SCM cultured bystander cells treated with the SIRT inhibitor. Excitingly, SIRT inhibition increased cholesterol (**Fig. 4G**). Consistent with the notion that SIRT activity is downstream of NDUFA5, SIRT inhibition, which does not affect detachment of bystander NDUFA5 knockdown cells cultured in SCM (**Fig. 4B**), also had no effect on nuclear SREBP1 (**Fig. 4E**) or cholesterol (**Fig. 4H**). Together, these data support the idea that the cisplatin-induced SAS promotes bystander cell detachment via NAD^+^-dependent SIRT activity downstream of NDUFA5, which inhibits SREBP1-medi-ated cholesterol synthesis (**Fig. 4I**).

**Figure 4.**
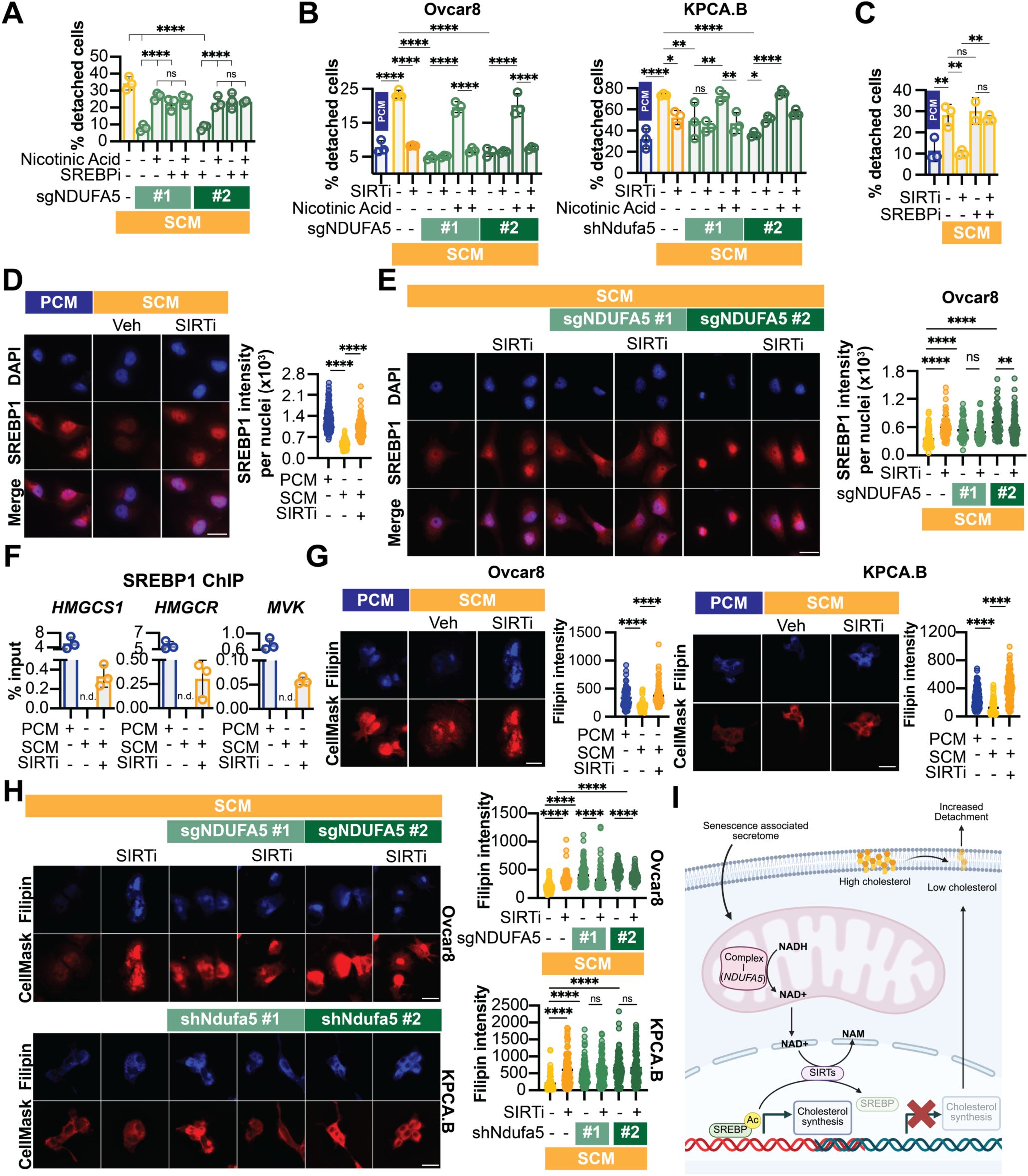
SREBP1 activity is decreased by the cisplatin-induced SAS downstream of NUDFA5 through NAD^+^-dependent SIRT activity. **(A)** Ovcar8 bystander cells with knockout of NDUFA5 (sgNDUFA5 #1 and #2) were cultured in SCM alone or treated with or without the SREBP inhibitor (SREBPi) fatostatin and/or nicotinic acid. After 72h the controlled trypsin experiment was performed. Shown are % detached cells. **(B)** The indicated bystander cells with knockout of NDUFA5 (sgNDUFA5 #1 and #2) or knockdown of Ndufa5 (shNdufa5 #1 and #2) were cultured in PCM or SCM alone and treated with or without a pan-SIRT inhibitor (SIRTi) with or without nicotinic acid. After 72h the controlled trypsin experiment was performed. Shown are % detached cells. **(C)** Ovcar8 bystander cells were cultured in PCM or SCM alone or treated with or without a pan-SIRT inhibitor (SIRTi) and/or the SREBP inhibitor (SREBPi) fatostatin. After 72h the controlled trypsin experiment was performed. Shown are % detached cells. **(D)** Ovcar8 bystander cells were cultured in PCM or SCM alone or treated with or without a pan-SIRT inhibitor (SIRTi). After 72h, cells were stained with anti-SREBP1 antibody and DAPI and imaged. Left: Representative images. Scale bar = 25 µm. 40X magnification. Right: Quantification of SREBP1 immunofluorescence intensity per nuclei. **(E)** Ovcar8 bystander cells with knockout of NDUFA5 (sgNDUFA5 #1 and #2) were cultured in SCM alone or treated with or without a pan-SIRT inhibitor (SIRTi). After 72h, cells were stained with anti-SREBP1 antibody and DAPI and imaged. Left: Representative images. Scale bar = 25 µm. 40X magnification. Right: Quantification of SREBP1 immunofluorescence intensity per nuclei. Representative data from 2 independent experiments (n >100 cells). **(F)** Ovcar8 bystander cells were cultured in PCM or SCM with or without a pan-SIRT inhibitor (SIRTi). After 72h, cells were harvested for chromatin immunoprecipitation (ChIP) using an anti-SREBP1 antibody following by qPCR. Shown is occupancy of SREBP1 in the promoter of *de novo* cholesterol synthesis genes *HMGCS1*, *HMGCR*, and *MVK*. n.d. = not detected. **(G)** The indicated bystander cells were cultured in PCM or SCM with or without a pan-SIRT inhibitor (SIRTi). After 72h, cholesterol was assessed using filipin III fluorescent staining. Left: Representative images. Scale bar = 25 µm. 20X magnification. Right: Quantification. n >100 cells. **(H)** The indicated bystander cells with knockout of NDUFA5 (sgNDUFA5 #1 and #2) or knockdown of Ndufa5 (shNdufa5 #1 and #2) were cultured in SCM alone or treated with or without a pan-SIRT inhibitor (SIRTi). After 72h, cholesterol was assessed using filipin III fluorescent staining. Left: Representative images. Scale bar: 20X magnification. Right: Quantification. **(I)** Schematic showing regulation of cholesterol synthesis downstream of SAS-mediated complex I activity though the NAD^+^-SIRT-SREBP axis. Panels B and H are representative data from 2 independent experiments in each cell line (B: n = 3; H: n>100 cells). Panels A, C, D, F, G, are representative data from 3 independent experiments (n=3). All graphs show mean ± SD; one-way ANOVA. ns = not significant, *p>0.05, *p<0.05, *p<0.01, ***p<0.001, ****p<0.0001.

### Increased fructose in the cisplatin-induced senescence-associated secretome drives the paracrine increase in cell detachment

Finally, we aimed to determine the upstream signal in the SAS that regulates detachment in a paracrine fashion. We found that both heat inactivated or <3kD-filtered SCM increase detachment (**Fig. 2E**), indicating that the signal promoting detachment is a small molecule or metabolite. Senescent cells are known to increase glucose uptake^21,26,75,76^, resulting in a glucose-depleted SAS. Indeed, we observed a significant decrease in glucose abundance in conditioned media from cisplatin-induced senescent cells versus vehicle controls (**Fig. 5A**). To understand how decreased glucose in the SAS affects bystander cells in a paracrine fashion, we performed mass spectrometry and assessed glycolytic intermediates. Unexpectedly, although conditioned media from cisplatin-induced senescent cells has decreased glucose (**Fig. 5A**), we observed an increase in glycolytic intermediates in bystander cells cultured in SCM (**Fig. S5A-B**). One notable metabolite that was upregulated in bystander cells cultured in SCM is dihydroxyacetone phosphate (DHAP) (**Fig. S5C**), which can be derived from fructolysis^77^ (**Fig. S5A**). Indeed, we found that conditioned media from cisplatin-induced senescent cells had increased fructose abundance (**Fig. 5B**), consistent with prior reports demonstrating that high glucose can induce polyol pathway activity, which converts glucose to fructose^78–80^. Remarkably, and in agreement with our findings, we found that elevated fructose is associated with higher stage at diagnosis in HGSOC patient, indicating more widespread disease (**Fig. 5C**). To determine how the relative availability of fructose and glucose contribute to bystander cell adhesion, we cultured bystander cells in PCM supplemented with fructose or SCM supplemented with glucose. The addition of fructose to PCM cultured bystander cells was sufficient to induce detachment in controls but not NDUFA5 knockdowns (**Fig. 5D**), demonstrating that fructose is upstream of NDUFA5. Conversely, the addition of glucose inhibited detachment of bystander cells cultured in SCM (**Fig. S5D**), indicating that the balance between fructose and glucose drives the observed detachment. Finally, we sought to determine whether fructose is upstream of decreased SREBP1 activity and cholesterol abundance. Addition of fructose to PCM, but not to NDUFA5 knockdown cells, was sufficient to decrease nuclear SREBP1 and cholesterol content of bystander cells (**Fig. 5E-H and Fig. S5E-F**). Conversely, the addition of glucose increased SREBP1 nuclear localization and cholesterol content of bystander cells cultured in SCM (**Fig. S5G-J**). Together, these data demonstrate that fructose suppresses nuclear SREBP1, corresponding to less plasma membrane cholesterol and increased detachment. These data also show that restoring the balance between glucose and fructose overcomes this phenotype in SCM cultured bystander cells.

**Figure 5.**
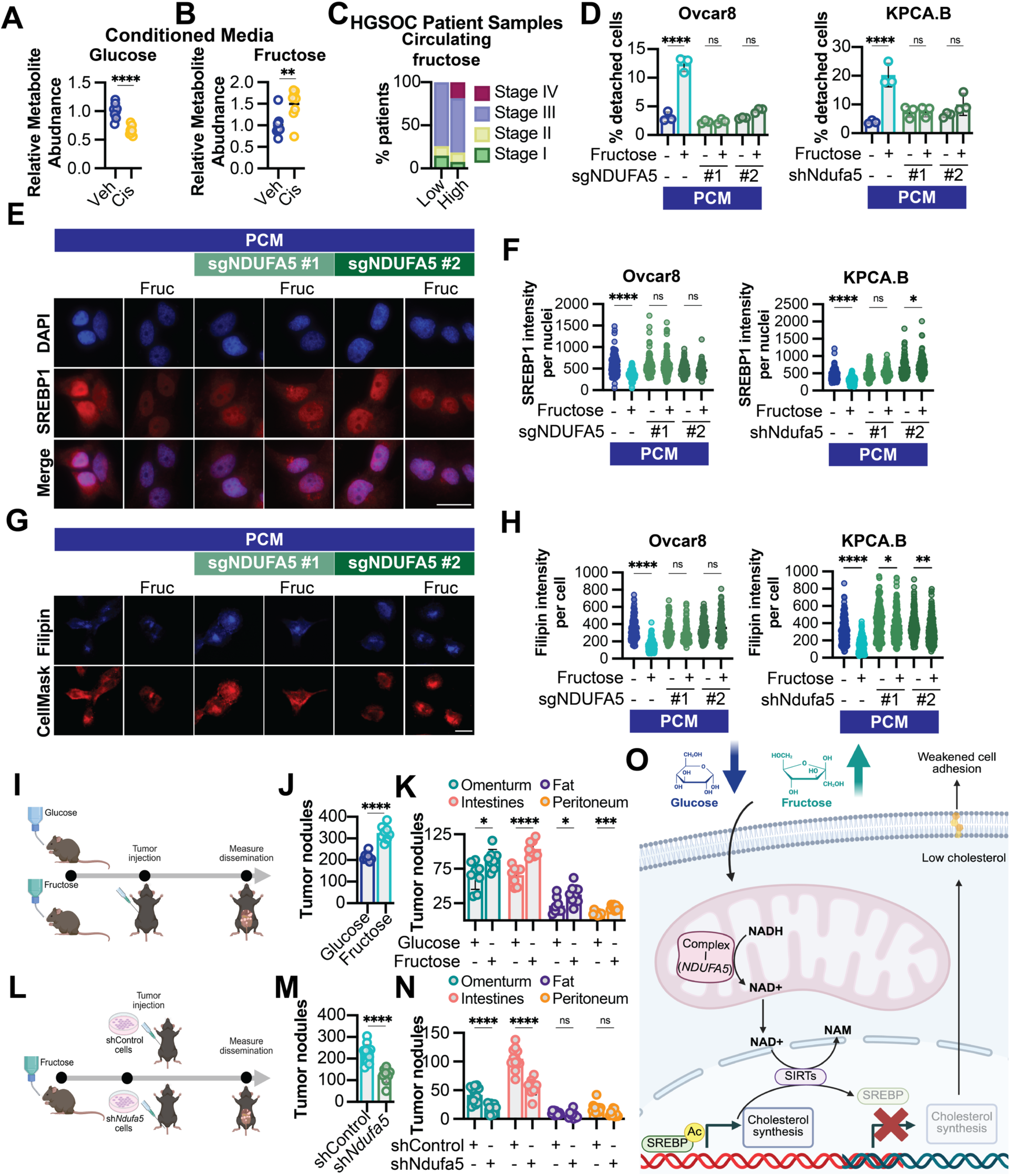
Fructose is increased in the cisplatin-associated senescent secretome and promotes HGSOC detachment and dissemination. (A-B) KPCA.B cells were induced to senesce using cisplatin, and conditioned media was harvested. **(A)** Relative glucose abundance normalized to cell AUC was assessed by LC-MS. Student’s t-test. **(B)** Relative fructose abundance normalized to cell AUC was assessed by LC-MS. Student’s t-test. **(C)** HGSOC patients from a University of Pittsburgh cohort were stratified by median circulating fructose. Shown is stage at diagnosis. **(D)** The indicated bystander cells with knockout of NDUFA5 (sgNDUFA5 #1 and #2) or knockdown of Ndufa5 (shNdufa5 #1 and #2) were cultured in PCM alone or supplemented with fructose. After 72h the controlled trypsin experiment was performed. Shown are % detached cells. One-way ANOVA. **(E-F)** The indicated bystander cells with knockout of NDUFA5 (sgNDUFA5 #1 and #2) or knockdown of Ndufa5 (shNdufa5 #1 and #2) were cultured in PCM alone or supplemented with fructose. **(E)** After 72h, cells were stained with anti-SREBP1 antibody and DAPI and imaged. Scale bar = 25 µm. 40X magnification. **(F)** Quantification of **(E)**. n >100 cells; one-way ANOVA. **(G-H)** The indicated bystander cells with knockout of NDUFA5 (sgNDUFA5 #1 and #2) or knockdown of Ndufa5 (shNdufa5 #1 and #2) were cultured in PCM alone or supplemented with fructose. **(G)** After 72h, cells were stained with filipin III to assess cholesterol and images. Scale bar = 25 µm. 40X magnification. **(H)** Quantification of **(G)**. n >100 cells; one-way ANOVA. **(I)** Schematic of *in vivo* experiment with mice provided fructose or glucose in the drinking water. **(J)** Endpoint total intraperitoneal tumor nodules from the experiment detailed in **(I)**. Student’s t-test. 8 mice/group. **(K)** Endpoint tumor nodule counts in the indicated metastatic sites from the experiment detailed in **(I)**. Student’s t-test. 8 mice/group. **(L)** Schematic of *in vivo* experiment with mice provided fructose or glucose in the drinking water and implanted with control (shControl) or Ndufa5 knockdown (shNdufa5) cells. **(M)** End-point tumor nodule counts in the indicated metastatic sites from the experiment detailed in **(L)**. Student’s t-test. 9-12 mice/group. **(N)** Endpoint tumor nodule counts in the indicated metastatic sites from the experiment detailed in **(L)**. Student’s t-test. 9-12 mice/group. **(O)** Final proposed model. High fructose in the SAS leads to metabolic reprogramming in bystander cells through the complex I-NAD^+^-SIRT-SREBP1 axis leading to low plasma membrane cholesterol and increased detachment. Panels A-B are representative data from 2 independent experiments in each cell line (n=8). Panels C-G are representative data from 3 independent experiments (n=3). All graphs show mean ± SD; ns = not significant, *p>0.05, *p<0.05, *p<0.01, ***p<0.001, ****p<0.0001.

We found that fructose is a component of the SAS and drives differential cell detachment *in vitro* (**Fig. 5B, 5D**). We also found that SCM increases HGSOC dissemination *in vivo* (**Fig. 1**). Thus, we interrogated whether fructose is sufficient to increase dissemination *in vivo*, independent of senescent cells or other SAS components. Towards this goal, we provided mice with fructose in their drinking water and implanted HGSOC cells (**Fig. 5I**). Glucose was used to control for water sweetness (**Fig. 5I**). Notably, we did not observe differences in water consumption between groups (**Fig. S5K**). High dietary fructose in-creased HGOSC dissemination in mice with similar tropism to mice co-injected with senescent cells (**Fig. 5J-K**) without differences in omental weights (**Fig. S5L**). Fructose can have pleotropic systemic effects^81^. Thus, we aimed to assess whether the increased fruc-tose-driven dissemination is due in part to cancer cell complex I activity. To this end, we provided mice with fructose and implanted control or shNdufa5 HGSOC cells (**Fig. 5L**). Ndufa5 knockdown abrogated dissemination in mice on a high fructose diet (**Fig. 5M-N**) but did not affect omental weights (**Fig. S5M**), which is consistent with our prior experiment using Ndufa5 knockdown cells co-injected with senescent cells (**Fig. 2M-O**). These data demonstrate the sufficiency of fructose to drive HGSOC dissemination through complex I activity *in vivo*. Together with mechanistic and patient data, these studies point to a previously unrecognized role for fructose in promoting cell detachment through a complex I-NAD^+^-SIRT-SREBP1-cholesterol synthesis axis and suggest that lowering dietary fructose may have beneficial clinical outcomes for HGSOC patients (**Fig. 5O**).

## DISCUSSION

The phenomenon of senescence has gained increasing attention in cancer research due to its complex role in influencing the TME through its unique secretome. In this study, we investigated the impact of TIS cell metabolism and the SAS on HGSOC progression to better understand how chemotherapy-induced metabolic reprogramming affects tumor progression. Our results demonstrate for the first time that the SAS from cisplatin-induced senescent HGSOC cells is sufficient to increase cell detachment and drive increased metastatic dissemination in HGSOC. Mechanistically, we found that the increased detachment is regulated by complex I, which leads to a reduction of cholesterol in the plasma membrane via SIRT-mediated SREBP1 activity, leading to increased detachment. Finally, we identified that the upstream metabolic signal in the SAS is fructose. These studies have significant implications for understanding how chemotherapy alters tumor dynamics in a paracrine fashion, unveil a novel mechanism of how senescence impacts bystander cancer cells on a molecular level, and reveal a potential role for diet in modifying HGSOC progression.

We found that the SAS from cisplatin-induced senescent HGSOC cells alone increases detachment of bystander cells and drive metastatic dissemination in a model of HGSOC. While prior studies have shown that senescent cells promote tumor progression and metastasis through co-injection or inducible models^9,10,82,83^, these effects reflect the combined influence of senescent cells and their secretome. In our model, by injecting only conditioned media from senescent cancer cells, we specifically isolate the impact of the SAS in the absence of senescent cells, demonstrating that secreted factors alone are sufficient to drive the metastatic phenotype (**Fig. 1**). While components of the SAS have been linked to metastasis in various cell and cancer types, many of these are either pro-inflammatory mediators such as hepatocyte growth factor, IL-6, and IL-8 or matrix remod-elers such as MMPs^11,13,84,85^. Unexpectedly, we found that the increase in SAS-mediated cell detachment was likely not due to MMP activity or other proteins (**Fig. 2C and 2E**). The non-protein, non-cytokine fractions of the SAS are often overlooked. In fact, there are only a handful of reports on the metabolic components of the SAS, such as elevated bioactive lipid production, nucleobases, amino acids, nitric oxide and ROS^21–23,86–88^. Moreover, only a small number of studies to date have indicated that specific metabolites are depleted from the SAS^21,26^. Notably, while these reports investigated the metabolite’s role in senescence induction, no study to date has investigated the paracrine effects of the extracellular metabolite composition on proliferative bystander cells. Here we found that glucose is depleted in the therapy-induced SAS (**Fig. 5A**). This is likely due to increased glucose uptake and glycolysis of senescent cells, which has been previously reported^21,26^. High glucose can induce polyol pathway activity, which converts glucose to fructose^78–80^. Indeed, we observed an increase in fructose in the SAS along with the concomitant decrease in glucose (**Fig. 5A-B**). Fructose production in malignant cells has been shown to increase respiration^89^ and also enhance proliferation and migration^90^, although none of this has been done in the context of senescence or chemotherapy. These data highlight the need to further characterize metabolites in the SAS and reinforce the role of fructose in pro-tumorigenic phenotypes.

We found that high dietary fructose increases metastatic dissemination *in vivo* (**Fig. 5**). While glucose is the most abundant monosaccharide in the body, fructose makes up a significant portion of sugar consumed. Fructose is present alone and together with glucose in sucrose in foods like honey, fruits, vegetables, and high-fructose corn syrup. Resting plasma fructose levels are in the tens of micromolar but can increase 10-fold after consuming fructose^91^. Fructose, or beverages with high fructose, have been associated with increased cancer risk and mortality in pancreatic, colorectal, breast, endometrial, and ovarian cancers^92–95^. Our data demonstrate that fructose has direct effects on cancer cells (**Fig. 5**). Exogenous fructose is absorbed passively in the gut through the transporter GLUT5^91^. GLUT5 has also been reported to be highly expressed in ovarian cancer cells, and its expression can support tumor growth^96^. Although our data implicate a role of fructose acting directly on malignant bystander cells, fructose metabolism can indirectly support tumor growth through the microbiome or liver lipogenesis^97,98^, and future studies will determine the contribution of these pathways to metastatic dissemination. Given the dietary and endogenous availability, and our findings on fructose driving dissemination, fructose may play an underappreciated role in modulating tumor cell behavior. This raises the possibility that restricting fructose intake could be a promising dietary intervention to assist in treatment response.

Our data reveal a critical role for respiration in reducing cell adhesion, allowing cells to detach and disseminate (**Fig. 2**). There is growing evidence that mitochondrial function and efficient respiration supports tumor progression and metastasis. In leukemia, high mitochondrial metabolism and mass are associated with relapse and increased oxidative phosphorylation is associated with therapy resistance^99,100^. ETC activity is also implicated in promoting relapse in pancreatic ductal carcinoma^101^. Complex I inhibition in particular can prevent melanoma brain metastasis formation^33^ and metastatic potential in response to low glucose^102^. Similarly, clear cell renal cell carcinomas, which have been show low glucose uptake, depend on complex I activity for metastasis driven by recycling of NADH to NAD^+31^. Likewise, our data supports the notion that plasma membrane cholesterol depletion in cancer cells can lead to a more disseminated phenotype (**Fig. 3**), raising questions about the use of statins in cancer patients. Interestingly, while long-term use of statins is context dependent in terms of altering cancer risk^103^, statins have had generally positive effects as a cancer intervention^104–106^. In contrast, recent mechanistic investigation into statin use in cancer revealed heterogenous responses, including increasing metastatic potential^107^. Taken together with our data, this highlights the potential context dependency in cholesterol metabolism in malignant cell phenotypes, though the interpretation of the systemic effects of long-term statin treatment can be difficult. The translation of these findings to ovarian cancer could represent a novel therapeutic strategy to specifically target senescent cells and mitigate the metastatic potential induced by the SAS.

We found that plasma membrane cholesterol is critical for SAS-mediated detachment through a SIRT- and SREBP1-dependent signaling axis (**Fig. 3-4**). Plasma membrane cholesterol content is important for plasma membrane structure, function, and signaling^108–111^, including to activate cell adhesion molecules such as integrins^112,113^. While the role of integrins and their stabilization by cholesterol in our model is unknown, the role of integrins in the metastatic processes in ovarian cancer has been previously discussed^54^; however, this is frequently in the context of how spheroids survive and invade after detachment from the primary solid tumor. Prior work has demonstrated that integrin expression and function promote spheroid formation^114^, and integrins are necessary for disseminating spheroids to adhere to a secondary site^115^. Additionally, dense compaction of spheroids enhances tumorigenesis as well as the invasive capacity of disseminating ovarian cancer spheroids^42,116^. Studies have also investigated the role of senescent mesothelial cells in promoting mesothelial clearance at secondary sites^117–119^. In this study, we aimed to understand the initial step in the metastatic dissemination cascade, which is likely biologically uncoupled from spheroid formation and adherence to metastatic sites. Regardless of the mechanism, cell adhesion has to be weakened in the initial detachment phase of dissemination^43–46^. Conversely, for cells to survive and adhere to metastatic sites, adhesion molecules and strong adhesion phenotypes are required^114,115,120^. Thus, the prior literature together with our data show the plasticity HGSOC cells display in order to disseminate and opens new questions about the dynamics of cell adhesion and detachment across different phases of dissemination.

It is interesting to speculate whether these changes in adhesion and detachment observed in bystander cells exposed to SCM could be therapeutically targeted. One such therapeutic route is the use of senolytics, a class of drugs designed to selectively target and eliminate senescent cells^121^. In the context of cancer, selective killing of senescent cells has exhibited anti-tumor effects, including suppressing metastasis^9,122^. Importantly, the senolytics dasatinib and quercetin have shown to reduce metastasis when combined with carboplatin or olaparib to treat ovarian cancer^122^. The potential benefits are underscored by their success in cancer types other than ovarian cancer. In breast cancer, GL-V9 has been shown to preferentially kill both senescent breast cancer cells and repli-cation-induced senescent fibroblasts^123^, and its use has been shown to decrease invasion and metastasis^124^. Similarly, the senolytic Digitonin has been shown to be an effective complement to chemotherapy in breast cancer treatment^125^. Given our dietary studies (**Fig. 5**), it is also interesting to speculate that diet interventions could be used in concert with senolytics to attain a greater benefit. This will be an area for the field to explore further. Finally, there is a large body of literature regarding the aging-related decline in NAD^+^ and surrounding the use of supplements in aging to replenish NAD^+^ levels ^126^. We found that supplementing cells with nicotinic acid increased detachment *in vitro* (**Fig. 2Q**), and prior work found that NAD^+^ promotes the pro-inflammatory SASP^127^. Thus, our studies herein would indicate that nicotinic acid supplements in cancer patients, regardless of age, may be detrimental, although this needs to be experimentally investigated.

In summary, this study sheds light on a previously unexplored facet of ovarian cancer by revealing how the secretome of cisplatin-induced senescent cells contributes to HGSOC phenotypes. The SAS-induced dissemination, metabolic reprogramming leading to weakened adhesion and increased detachment, and the metabolite driving these changes provide a novel perspective on the complex interplay between senescence and cancer progression. These findings underscore the importance of understanding all components of the SAS and its multifaceted impact on the tumor microenvironment, offering potential therapeutic targets for mitigating ovarian cancer metastasis.

## ACKNOWELDGEMENTS

We would like to thank Uma Chandran and Jiefei Wang (UPMC Hillman Cancer Center Bioinformatics Core) for help with bioinformatic analysis of the CRISPR screen, and Maureen Lyons (UPMC Hillman Cancer Center Cancer Genomics Core) for help with sequencing of the CRISPR library. Multiple schematics were created with BioRender.com. This work was supported by National Institutes of Health (R37CA240625 to KMA, R01CA259111 to KMA and NWS, R01CA298386 to KMA and NWS, P50CA272218, T32GM133332 to ARC, R01CA242021 to NH, R21CA267050 to FM, R21CA291905 to FM, U01AG077923), the American Cancer Society (RSG-19-113-01-CCG to KMA), the Ovarian Cancer Research Alliance (MIG-2023-2-1018 to AU), Congressionally Directed Medical Research Program (HT9425-23-1-0436 to KMA, W81XWH2110338 to FM, OC210139 to LGC, OC230324 to NKT), HERA Ovarian Cancer Foundation (to NKT, AU, and AA), the Melanoma Research Foundation (to RB), the Janet Burroughs Ovarian Cancer Foundation (to FM), 2023-329680 from the Chan Zuckerberg Initiative DAF, an advised fund of Silicon Valley Community Foundation (to KK), and the UPMC Hillman Cancer Center. This project used the Hillman Animal Facility, Cancer Genomics Facility, Tissue and Research Pathology/Pitt Biospecimen Core, Biostatistics Core, and the Cancer Bioinformatics Services that are supported in part by award P30CA047904.

## Author Contributions

Aidan R. Cole: Conceptualization, Investigation, Methodology, Visualization, Writing – Original Draft, Visualization, Writing – Review & Editing, Funding Acquisition. Raquel Buj: Investigation, Methodology, Writing – Review & Editing. Apoorva Uboveja: Investigation, Methodology, Writing – Review & Editing. Evan Levasseur: Investigation. Hui Wang: Investigation. Katarzyna M. Kedziora: Investigation, Methodology. Adam Chatoff: Investigation, Methodology. Andrea Andress Huacachino: Investigation, Methodology. Mariola M. Marcinkiewicz: Investigation, Methodology. Amandine Amalric: Investigation, Methodology, Writing – Review & Editing. Baixue Yang: Investigation, Writing – Review & Editing. Naveen Kumar Tangudu: Investigation, Writing – Review & Editing. Jeff Danielson: Investigation. Amal Elwah: Investigation, Methodology. Sierra White: Investigation, Methodology. Danyang Li: Investigation, Methodology. Callen T. Wallace: Investigation, Methodology. Felicia Lazure: Investigation. Esther Elishaev: Resources. Lauren Borho: Resources. Dorota E. Jazwinska: Investigation, Methodology. Matthew S. Laird: Investigation, Methodology. Huda Atiya: Methodology. Benjamin G. Bitler: Writing – Review & Editing. Denarda Dangaj: Investigation, Methodology. Lan G. Coffman: Methodology, Supervision. George Tseng: Supervision. Steffi Oesterreich: Methodology, Writing – Review & Editing. Ana P. Gomes: Supervision, Writing – Review & Editing. Aditi U. Gurkar: Methodology, Supervision, Writing – Review & Editing. Francisco J. Schopfer: Methodology, Supervision, Writing – Review & Editing. Francesmary Modugno: Resources, Supervision. Simon C. Watkins: Investigation, Methodology, Supervision. Ioannis K. Zervantonakis: Investigation, Methodology, Supervision. Wayne Stallaert: Investigation, Methodology, Supervision. Nadine Hempel: Investigation, Methodology, Supervision, Writing – Review & Editing. Nathaniel W. Snyder: Investigation, Methodology, Supervision, Writing – Review & Editing. Katherine M. Aird: Conceptualization, Visualization, Writing – Original Draft, Writing – Review & Editing, Supervision, Project Administration, Funding Acquisition.

## Declaration of Interests

All authors declare no competing interests.

## SUPPLEMENTAL FIGURES

**Figure S1.**
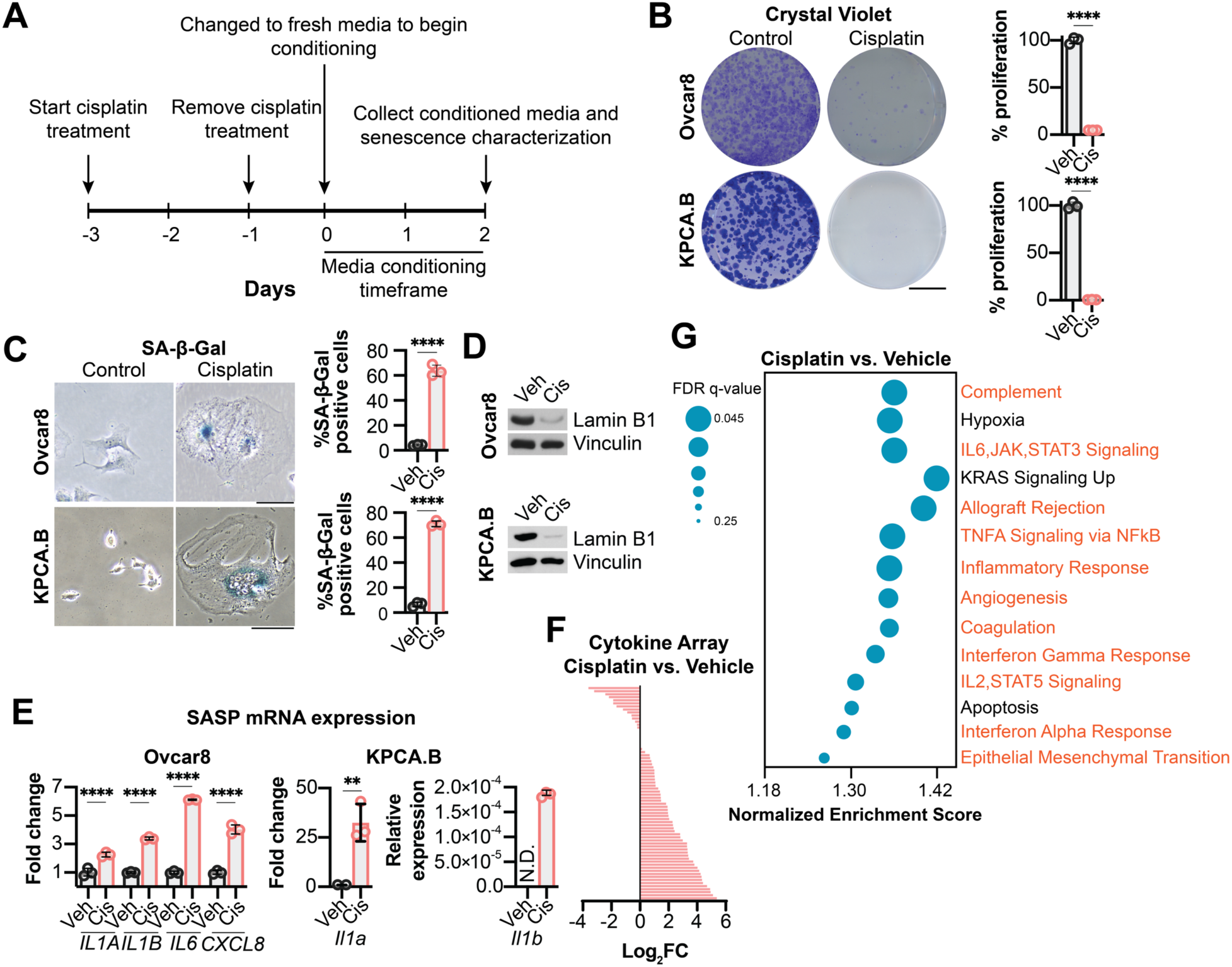
Confirmation of cisplatin-induced senescence in HGSOC models. Related to Figures 1-6. **(A)** Timeline of senescence induction experiments in HGSOC cells using cisplatin indicating when conditioned media was harvested and senescence in cells was characterized. **(B-E)** The indicated cells were treated with cisplatin as in Fig. S1A. **(B)** Cells were stained with crystal violet (left: representative images, scale bar = 10 mm.; right: quantification). **(C)** Senescence-associated-beta-galactosidase (SA-β-Gal) activity (left: representative images, scale bar = 10 µm, 20X magnification; right: quantification). **(D)** Western blot of Lamin B1. Vinculin was used as a loading control. **(E)** Relative expression of canonical SASP genes by qPCR. N.D. = not detected. **(F)** Secretion of cytokines by cytokine array from conditioned media of Ovcar8 cells treated with cisplatin. **(G)** GSEA analysis enriched pathways of RNA-Seq data from Ovcar8 cells treated with cisplatin. Pathways highlighted in red are related pathways associated with the SASP. Panels B-E are representative data from 3 independent experiments (n=3). Panels F-G are data from 1 experiments (n=2-3). All graphs show mean ± SD; Student’s t-test; **p<0.01, ****p<0.0001.

**Figure S2.**
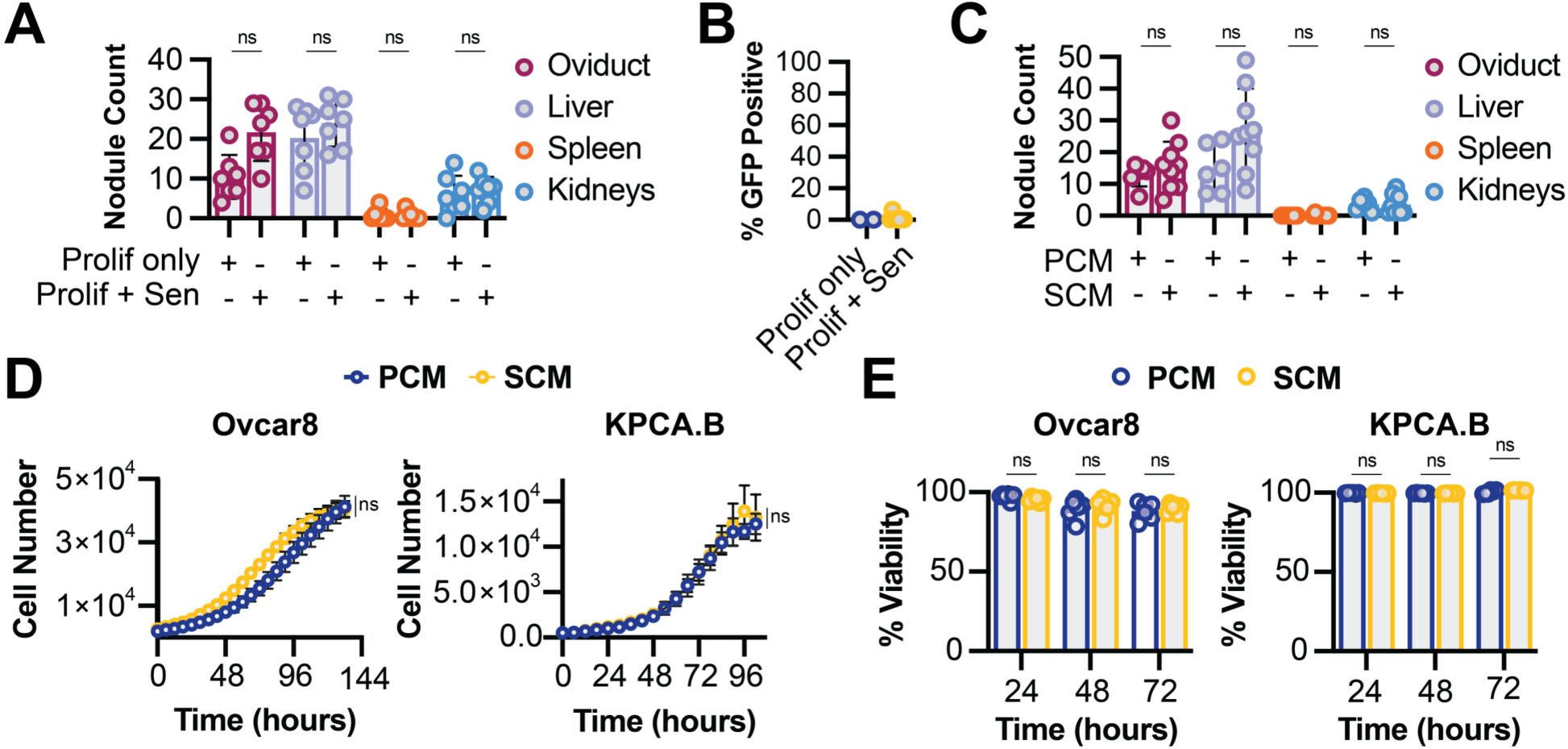
Some metastatic sites are not affected by the cisplatin-induced senescence-associated secretome; senescent cells do not gain proliferative ability *in vivo*; the cisplatin-induced senescence-associated secretome does not change proliferation or anoikis resistance in by-stander cells. Related to Figure 1. **(A)** KPCA.B HGSOC cells were treated with vehicle or induced to senesce using cisplatin. Cisplatin-induced senescent cells were also labeled with GFP. Vehicle treated or cisplatin-induced senescent cells were then co-injected intraperitoneally with naïve proliferating KPCA.B HGSOC. The same number of proliferating cells were injected into mice in each group, and the senescent cells were injected with naïve proliferating cells at a 1:1 ratio. Prolif only = vehicle treated cells co-injected with naïve proliferating cells. Prolif + Sen = cisplatin-induced senescent cells co-injected with naïve proliferating cells. Shown are endpoint tumor nodules in the indicated sites. 6-7 mice/group. **(B)** Percentage of total cells labeled with GFP recovered from disaggregated omental tumors from **(A)**. **(C)** KPCA.B HGSOC cells were treated with vehicle or induced to senesce using cisplatin, and conditioned media was harvested (see Methods for more details). PCM = proliferative conditioned media, which is conditioned media harvested from vehicle treated cells. SCM = senescent conditioned media, which is conditioned media harvested from cisplatin-induced senescent cells. KPCA.B HGSOC cells were injected intraperitoneally into mice, and upon tumor establishment, conditioned media was injected 3X per week. Shown are endpoint tumor nodules in the indicated sites. 6-9 mice/group. **(D)** The indicated bystander cells were cultured in PCM or SCM, and proliferation was assessed using Incucyte live cell imaging. **(E)** The indicated bystander cells in ultra-low attachment plates were cultured in PCM or SCM, and viability was assessed using fluorescent imaging of calcein AM (live cell probe) and ethidium homodimer (dead cell probe). Panels D-E are representative data from 3 independent experiments (n=6). All graphs show mean ± SD; Student’s t-test; ns = not significant.

**Figure S3.**
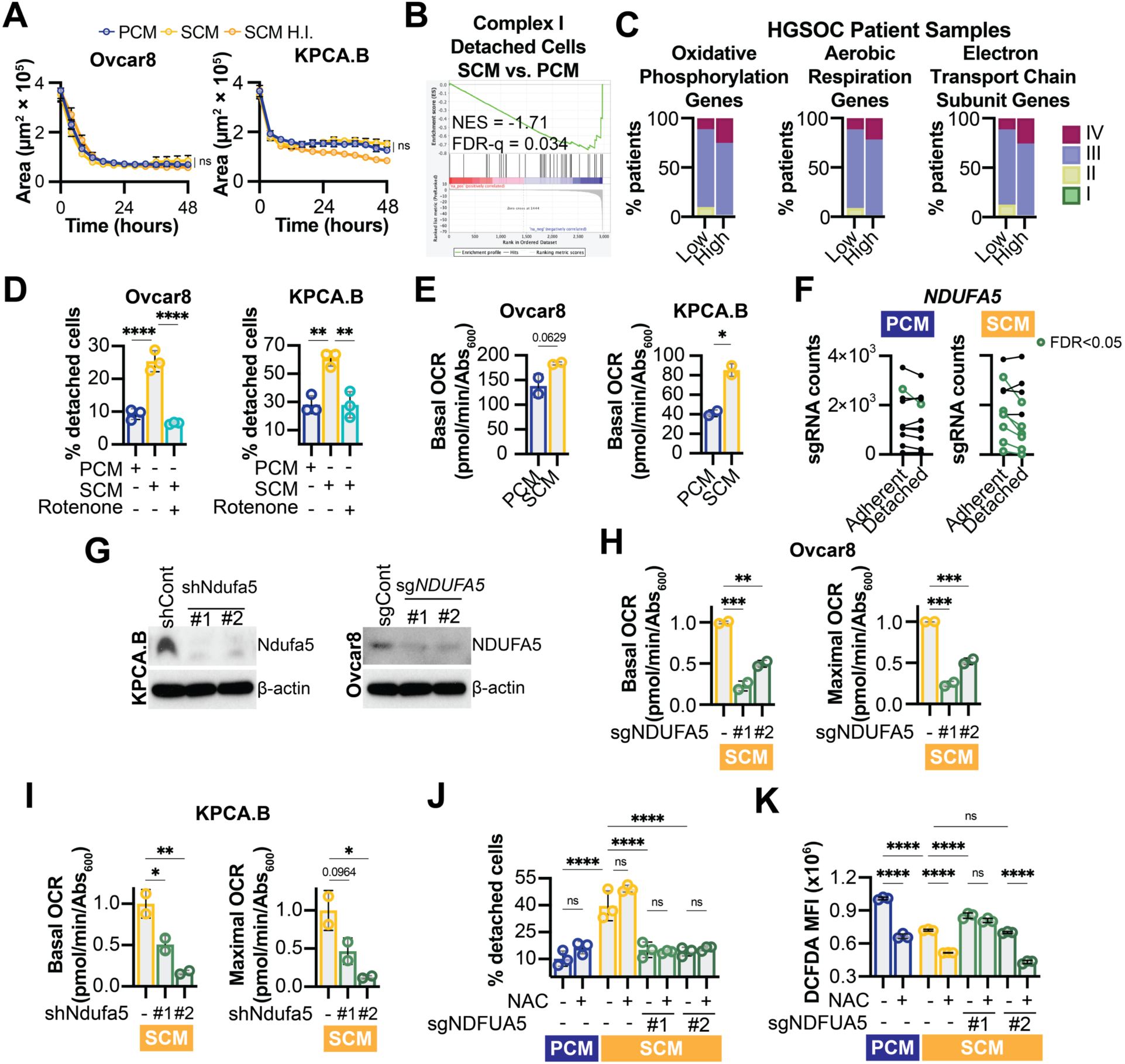
HGSOC bystander cells cultured in SCM do not show differences in compaction; HGSOC patients with high expression of oxidative phosphorylation, aerobic respiration, and electron transport chain genes have higher stage; NDUFA5 is required for increased detachment in SCM cultured bystander HGSOC cells, which is related to respiration but not reactive oxygen species. Related to Figure 2. **(A)** The indicated HGSOC bystander cells were cultured in proliferative conditioned media (PCM), senescent conditioned media (SCM), or heat inactivated SCM (SCM H.I.) in ultra-low attachment round bottom plates, and spheroid compaction was assessed using Incucyte live cell imaging. Data show mean ± SD. **(B)** Gene set enrichment analysis of the complex I submit gene set in detached cells cultured in SCM vs. PCM from the CRISPR screen. **(C)** HGSOC patients from TCGA were stratified based on the sum of expression of the indicated genes (see **Table S1** for gene lists). Shown is the stage of patients in the lower and upper quartile. **(D)** The indicated bystander cells were cultured in PCM or SCM with or without rotenone. After 72h, the controlled trypsin experiment was performed. Shown are % detached cells. Graph shows mean ± SD; one-way ANOVA. **(E)** The indicated bystander cells were cultured in PCM or SCM. After 72h, the basal respiration of cells was assessed using a Seahorse instrument. Graph shows mean ± SD; Student’s t-test. **(F)** Individual sgRNAs targeting *NDUFA5* in the controlled trypsin CRISPR screen. **(G)** The indicated cells were transduced with sg/shRNA controls or sg/shRNA targeting NDUF5A. Western blot of NDUFA5 protein expression. β-actin was used as a loading control. **(H-I)** The indicated bystander cells (**H**-Ovcar8, **I**-KPCA.B) were cultured in SCM. After 72h, the basal and maximal respiration of cells was assessed using a Seahorse instrument. Data show mean ± SD; one-way ANOVA. **(J-K)** Ovcar8 bystander HGSOC cells with knockout of NDUFA5 (sgNDUFA5 #1 and #2) were cultured in PCM or SCM with or without supplementation with N-acetyl-cysteine (NAC). **(J)** After 72h, the controlled trypsin experiment was performed. Shown are % detached cells. Graphs show mean ± SD; one-way ANOVA. **(K)** After 72h, ROS were assessed using the fluorescent probe DCFDA. Graph show mean ± SD; one-way ANOVA. Panels A-B, D-E, G-K are representative data from 3 independent experiments (n=2-3). ns = not significant, *p<0.05, *p<0.01, ***p<0.001, ****p<0.0001.

**Figure S4.**
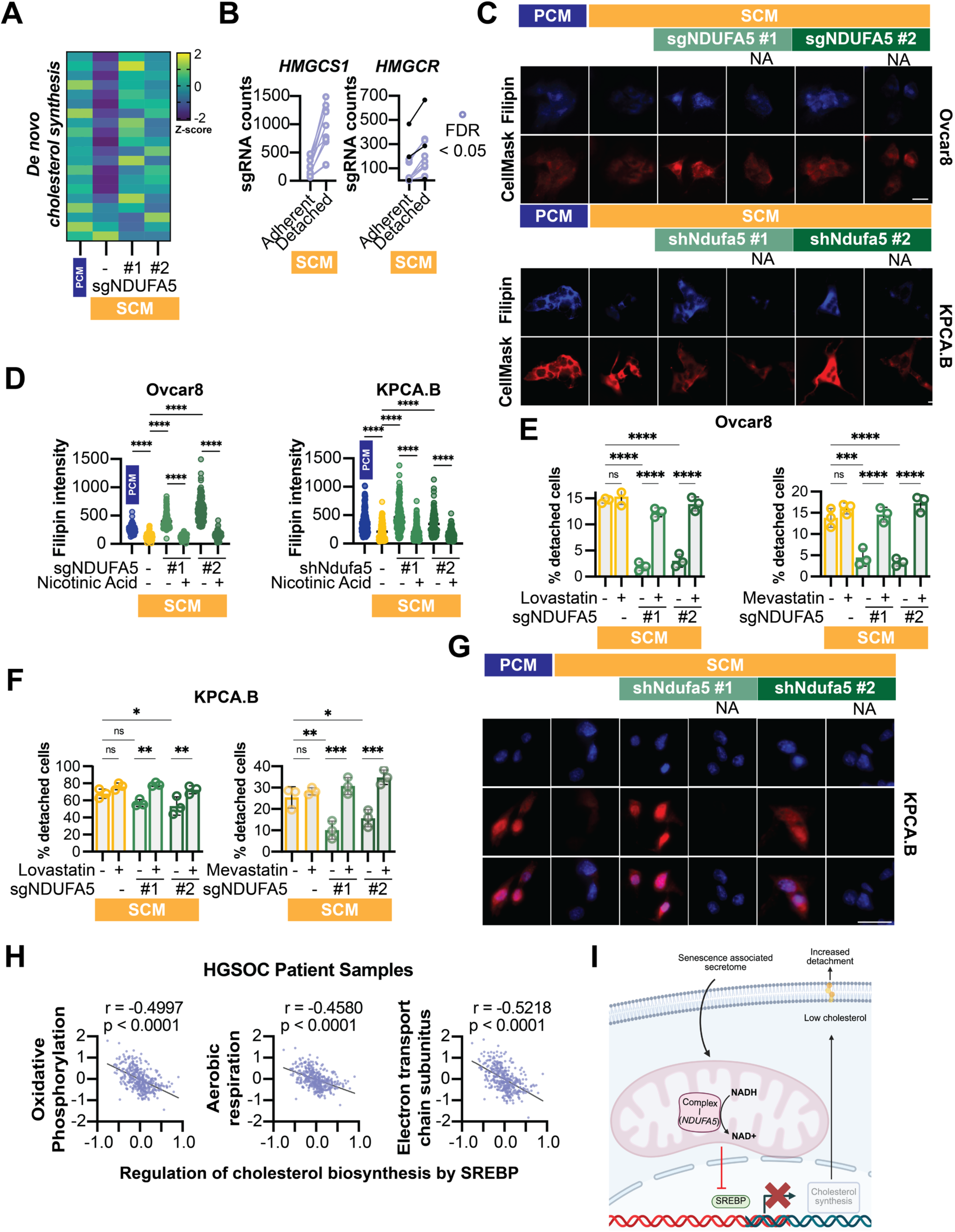
Bystander cells cultured in SCM have decreased expression of *de novo* cholesterol synthesis genes that is reversed by NDUFA5 knockout; detachment in SCM is decreased by supplementing with nicotinic acid and increased by inhibition of de novo cholesterol synthesis using statins; oxidative phosphorylation and associated genes negatively correlate with *de novo* cholesterol synthesis genes in HGSOC patient tumors. Related to Figure 3. **(A)** Ovcar8 bystander cells with either knockout of NDUFA5 (sgNDUFA5 #1 and #2) were cultured in PCM or SCM for 72h, and RNA was extracted. RNA-Seq was performed. Shown is a heatmap of *de* novo cholesterol genes. **(B)** Individual sgRNAs targeting HMGCS1 and HMGCR in the SCM cultured groups in the controlled trypsin CRISPR screen. **(C)** The indicated bystander cells with knockout of NDUFA5 (sgNDUFA5 #1 and #2) or knockdown of Ndufa5 (shNdufa5 #1 and #2) were cultured in PCM or SCM with or without nicotinic acid (NA). After 72h, total cholesterol was assess using filipin III fluorescent staining. Shown are representative images. Scale bar = 25 µm. 20X magnification. **(D)** Quantification of filipin III staining in **(C)**. **(E)** Ovcar8 bystander cells with knockout of NDUFA5 (sgNDUFA5 #1 and #2) were cultured in PCM or SCM with or without lovastatin or mevastatin. After 72h, the controlled trypsin experiment was performed. Shown are % detached cells. **(F)** KPCA.B bystander cells with knockdown of Ndufa5 (shNdufa5 #1 and #2) were cultured in PCM or SCM with or without lovastatin or mevastatin. After 72h, the controlled trypsin experiment was performed. Shown are % detached cells. **(G)** KPCA.B bystander cells with knockdown of Ndufa5 (shNdufa5 #1 and #2) were cultured in PCM or SCM with or without nicotinic acid. After 72h, total cholesterol was assess using filipin III fluorescent staining. Shown are representative images. Scale bar = 25 µm. 20X magnification. Quantification can be found in Fig. 3G. (H) Negative correlation between the expression of genes related with oxidative phosphorylation and cholesterol synthesis genes in TCGA HGSOC patient tumors (see **Table S1** for gene lists). Pearson’s correlation. **(I)** Schematic showing regulation of cholesterol synthesis through inhibition of SREBP downstream of complex I activity leading to low plasma membrane cholesterol and weakened adhesion. Panels C-F are representative data from 3 independent experiments (n=3). Graphs show mean ± SD; ns = not significant, *p>0.05, *p<0.05, *p<0.01, ***p<0.001, ****p<0.0001.

**Figure S5.**
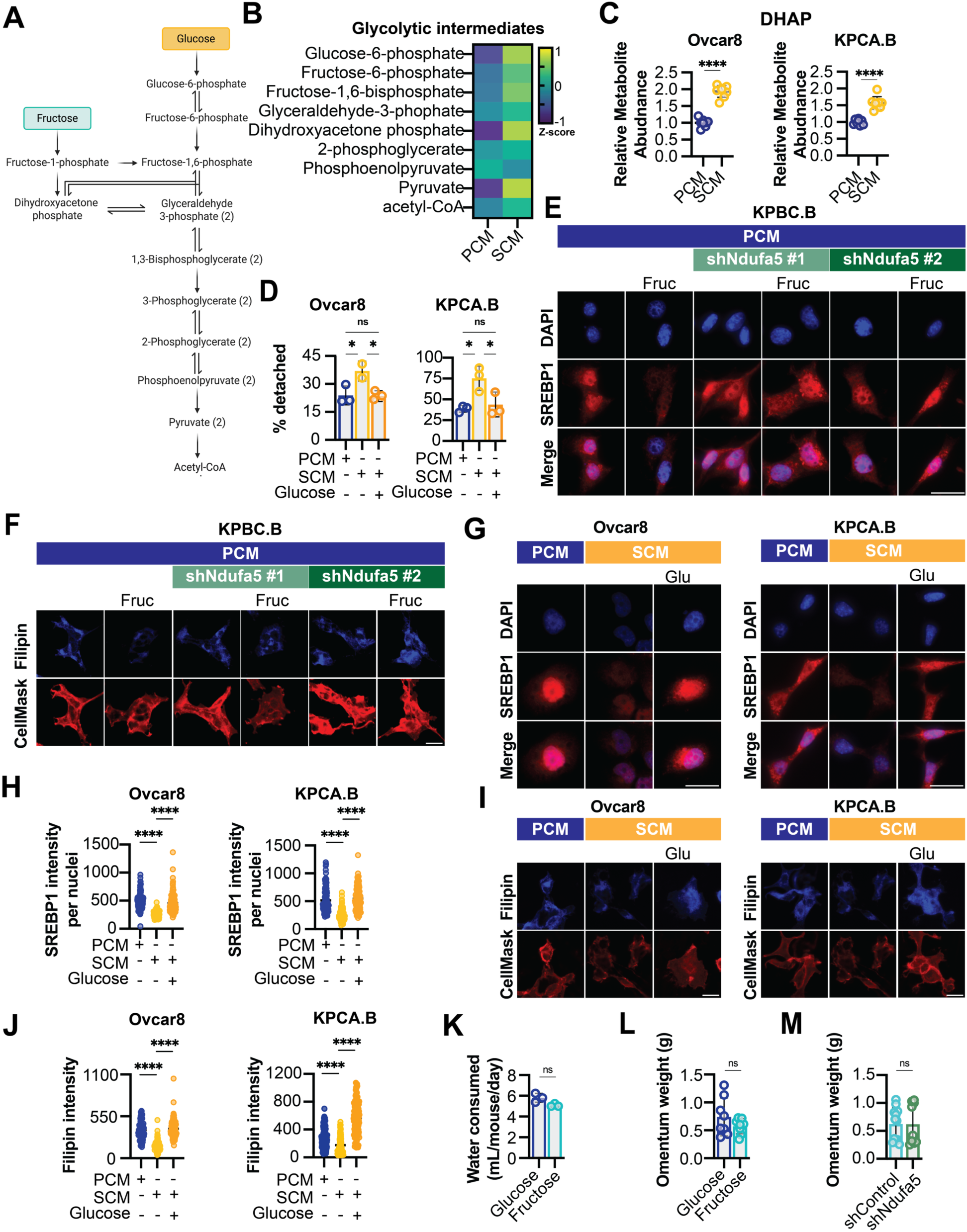
Glycolytic intermediates and DHAP are increased in bystander cells cultured in SCM; fructose decreases nuclear SREBP1 and cholesterol; glucose increases nuclear SREBP1 and cholesterol; mice on fructose or glucose water do not show differences in water consumption; no difference in omental weights at endpoint. Related to Figure 5. **(A)** Schematic of the fructolytic and glycolytic pathways. **(B)** Ovcar8 bystander cells were cultured in PCM or SCM. After 72h, cells were harvested and glycolytic intermediates were assessed using LC-MS. Shown is a heatmap of z-scores. Representative data from 1 independent experiment (n=8). **(C)** The indicated bystander cells were cultured in PCM or SCM. After 72h, cells were harvested. Relative DHAP abundance normalized to cell AUC was assessed by LC-MS. Student’s t-test. **(D)** The indicated bystander cells were cultured in PCM or SCM alone or supplemented with glucose. After 72h, the controlled trypsin experiment was performed. Shown are % detached cells. One-way ANOVA. **(E)** KPCA.B by-stander cells with knockdown of Ndufa5 (shNdufa5 #1 and #2) were cultured in PCM alone or supplemented with fructose. After 72h, cells were stained with anti-SREBP1 antibody and DAPI and imaged. Shown are representative images. Scale bar = 25 µm. 40X magnification. Quantification can be found in Fig. 5F. **(F)** KPCA.B bystander cells with knockdown of Ndufa5 (shNdufa5 #1 and #2) were cultured in PCM alone or supplemented with fructose. After 72h, cells were stained with filipin III and imaged. Shown are representative images. Scale bar = 25 µm. 40X magnification. Quantification can be found in Fig. 5H. **(G-H)** The indicated bystander cells were cultured in PCM or SCM alone or supplemented with glucose. **(G)** After 72h, cells were stained with anti-SREBP1 antibody and DAPI and imaged. Shown are representative images. Scale bar = 25 µm. 40X magnification. **(H)** Quantification of **(G)**. One-way ANOVA. Representative data from 2 independent experiments (n >100 cells). **(I-J)** The indicated bystander cells were cultured in PCM or SCM alone or supplemented with glucose. **(I)** After 72h, cells were stained with filipin III and imaged. Shown are representative images. Scale bar = 25 µm. 20X magnification. **(J)** Quantification of **(I)**. One-way ANOVA. Representative data from 2 independent experiments (n >100 cells). **(K)** Average water intake of mice provided glucose or fructose in their drinking water. **(L-M)** Omental weights from mice at endpoint. ns = not significant. Graphs show mean ± SD; 8 mice/group; Student’s t-test. Panels B-C are data from 1 experiment (n=8). Panels D-J are representative data from 3 independent experiments (n=3). Graphs show mean ± SD; ns = not significant, *p<0.05, ****p<0.0001.

## MATERIALS AND METHODS

### Cell Lines

Ovcar8 cells were a gift from Dr. Benjamin Bitler (University of Colorado) and cultured in RPMI 1640 (Fisher Scientific cat#) supplemented with 5% Fetal Bovine Serum (BioWest, cat# S1620) and 1% Penicillin/Streptomycin (Fisher Scientific, cat#15-140-122) unless otherwise noted. KPCA.B cells were a gift from Robert Weinberg and cultured in high glucose DMEM (Fisher Scientific cat#) supplemented with 4% Heat Inactivated Fetal Bovine Serum (Sigma, cat# S1620), 2 ng/mL hEGF (Sigma, cat# E9644), ITS-G (Fisher Scientific, cat# 41400-045) and 1% Penicillin/Streptomycin (Fisher Scientific, cat#15-140-122) unless otherwise noted. HEK293FT cells were used for lentiviral packaging and were cultured in DMEM (Corning, cat#10-013-CV) supplemented with 10% FBS according to ATCC. All cell lines were tested monthly for mycoplasma as described in ^128^.

### Senescence induction and conditioned media generation

Cells were treated with 1 µM (ovcar8) or 10 µM (KPCA.B) cisplatin (Selleck Chemicals, cat#S1166) or vehicle (DMF) for 48 hours, after which the cisplatin treated cells were washed with PBS and cultured in fresh media. Vehicle treated cells were split into fresh media. After 24 hours, the media was changed and allowed to condition for 48 hours. For conditioned media intended for use *in vivo*, no Penicillin/Streptomycin was added. Conditioned media was collected and filtered with a 0.22 mm sterile vacuum filter (Fisher Scientific, cat#FB12566506) and stored at −0° C. See the schematic in **Figure S1A**.

### Proliferation assays

For proliferation assays with crystal violet staining, an equal number of cells were seeded in multiwell plates and cultured for 7-10 days. Proliferation was assessed by fixing the plates for 5 min with 1% paraformaldehyde after which they were stained with 0.05% crystal violet. Wells were destained using 10% acetic acid. Absorbance (590nm) was measured using a spectrophotometer (BioTek Epoch Microplate reader). For proliferation assays were performed using an IncuCyte S3 imaging system (Sartorius), an equal number of cells per condition were seeded in triplicates in a 96-well plate. Cells were imaged for 4-5 days with live cell imaging every 6 hours. Cell quantification was performed using the IncuCyte software.

### Senescence Associated-β-Galactosidase (SA-β-Gal) assay

SA-β-Gal staining was performed as previously described ^129^. Cells were fixed in 2% formaldehyde/0.2% glutaraldehyde in PBS (5 min) and stained (40 mM Na_2_HPO_4_, 150 mM NaCl, 2 mM MgCl_2_, 5 mM K_3_Fe(CN)_6_, 5 mM K_4_Fe(CN)_6_, and 1 mg/ml X-gal) overnight at 37°C in a non-CO2 incubator. Images were acquired at room temperature using an inverted microscope (Nikon Eclipse Ts2) with a 20X/0.40 objective (Nikon LWD) equipped with a camera (Nikon DS-Fi3). Each sample was assessed in triplicate and at least 100 cells per well were counted (>300 cells per experiment).

### RNA isolation, Sequencing, and Analysis

Total RNA was extracted from cells with Trizol (Ambion, cat#15596018) and DNase treated, cleaned, and concentrated using Zymo columns (Zymo Research, cat#R1013) following manufacturer’s instructions. RNA integrity number (RIN) was measured using BioAnalyzer (Agilent Technologies) RNA 6000 Nano Kit to confirm RIN above 7 for each sample. The cDNA libraries, next generation sequencing, and bioinformatics analysis was performed by Novogene. Raw and processed RNA-Seq data can be found on GEO (GSE305033 and GSE305002).

### RT-qPCR

RNA was retrotranscribed with High-Capacity cDNA Reverse Transcription Kit (Applied Biosystems, cat#4368814) and 20ng of cDNA amplified using the CFX Connect Real-time PCR system (Bio-Rad) and the PowerUp^TM^ SYBR^TM^ Green Master Mix (Applied Biosystems, cat#A25742) following manufacturer’s instructions. Primers were designed using the Integrated DNA Technologies (IDT) web tool (**Table S7**). Conditions for amplification were: 5 min at 95° C, 40 cycles of 10 sec at 95° C and 7 sec at 62° C. The assay ended with a melting curve program: 15 sec at 95° C, 1 min at 70° C, then ramping to 95° C while continuously monitoring fluorescence. Each sample was assessed in triplicate. Relative quantification was determined to multiple reference genes (**Table S7**) to ensure reproducibility using the delta-delta CT method.

### Western blotting

Cells lysates were collected in 1X sample buffer (2% SDS, 10% glycerol, 0.01% bromophenol blue, 62.5mM Tris, pH 6.8, 0.1M DTT) and boiled to 95°C for 10 min. Protein concentration was determined using the Bradford assay (Bio-Rad, cat#5000006). An equal amount of total protein was resolved using SDS-PAGE gels and transferred to nitrocellulose membranes (Fisher Scientific) at 110mA for 2h at 4°C. Membranes were blocked with 5% nonfat milk or 4% BSA in TBS containing 0.1% Tween-20 (TBS-T) for 1h at room temperature. Membranes were incubated overnight at 4°C in primary antibodies in 4% BSA/TBS + 0.025% sodium azide. Membranes were washed 4 times in TBS-T for 5 min at room temperature after which they were incubated with HRP-conjugated secondary antibodies for 1 h at room temperature. After washing 4 times in TBS-T for 5 min at room temperature, proteins were visualized on film after incubation with SuperSignal West Pico PLUS Chemiluminescent Substrate (ThermoFisher, Waltham, MA).

### Cytokine array

The human cytokine antibody array C1000 (RayBio, cat# AAH-CYT-1000-2) was used to quantify secreted factors as we have previously published^130^. Conditioned medias were obtained as described above. Membranes were visualized on film. Individual spot signal densities were obtained using ImageJ software and normalized to cell number from which the conditioned media were obtained.

### Murine ovarian cancer *in vivo* models

Six-week-old female C57BL/6J mouse mice were purchased from Jackson Laboratories (cat# 2498079-86). All mice were maintained in a HEPA-filtered ventilated rack system at the Animal Facility of the Assembly Building of The Hillman Cancer Center at the University of Pittsburgh School of Medicine. Mice were housed up to 5 mice per cage and in a 12-hour light/dark cycle. All experiments with animals were performed in accordance with institutional guidelines approved by the Institutional Animal Care and Use Committee (IACUC) at the University of Pittsburgh School of Medicine (protocol ID: 20108084). For dietary studies with glucose and fructose, regular water was replaced with solutions of 30% w/v of sugar in sterile water 10 days prior to cell implantation and continued throughout the end of the experiment. One million KPCA.B ovarian cancer cells with or without *Ndufa5*^KD^ were injected intraperitoneally alone or together with one million cisplatin ther-apy-induced senescent parental KPCA.B cells. For conditioned media treatment experiments, mice were treated three times per week with intraperitoneal injection of 200 µL of conditioned media from proliferative or cisplatin therapy-induced senescent cells. Mice were imaged by IVIS at 5-8 days post injection and weighed weekly. Animals were euthanized at the end of fourth week, and tumor burden was assessed by counting the number of intraperitoneal nodules. TIF was harvested from omental tumors as previously described ^131^. Omentum tumors and intestinal metastasis were collected and either fresh frozen or formalin fixed and paraffin embedded.

### Immunohistochemistry (IHC)

IHC was performed with slight modifications from previously described methods ^132^. After deparaffinization and rehydration, tissues were steamed for 40min in 1X citrate buffer (Sigma-Aldrich, Cat#: C9999) after which they were immersed in 3% MeOH for 20min. Tissues were blocked with 1% BSA in PBS for 30min prior to overnight incubation in primary antibodies. Mouse and Rabbit Specific HRP/DAB (ABC) Detection IHC kit (Abcam, Cat#64264) was used for visualization of staining. Tissues were incubated in Mayer’s Hematoxylin Solution (Sigma-Aldrich, Cat#MHS16) for 3min followed by bluing in running tap water for 2min. Tissues were dehydrated and mounted in Cytoseal (Sigma-Aldrich, Cat# 23-244257). QuPath was used for H-Score analysis and to analyze the % of positive cells.

### Patient stage and transcriptomic data from TCGA

mRNA data were extracted from The Cancer Genome Atlas (TCGA) Serous Ovarian Cancer (Nature, 2011) project was downloaded from cbioportal.org. Analysis of tumor stage was performed by splitting patients into quartiles of the average z-score of the genes in the indicated gene sets. For correlative analysis, the average z-score of the genes in the indicated gene sets were plotted and linear regression was performed using GraphPad Prism 10. All gene sets can be found in **Table S1**.

### Non-adherent culture and imaging

Non-adherent cell culture and imaging was performed as previously described ^133^. Briefly, cells were seeded 1,000 (96 well round bottom ULA, Corning cat# 7007), 10,000 (96 well flat bottom ULA, Corning cat# 3474), or 300,000 (6 well flat bottom ULA, Corning cat# 3471) cells per well in a 1:1 ratio of fresh to conditioned media. To assess viability, cells were treated with 1 mM ethidium homodimer (Sigma Aldrich, cat# 46043) and 0.5 mM calcein AM (Invitrogen, cat#C1430) for 30 minutes and images were acquired using a Leica Thunder Imager. Timelapse imaging to assess spheroid formation in round bottom ULA plates was performed using IncuCyte S3 imaging system (Sartorius). Cells were seeded in 96 well ULA plates and images every 3 hours for 48 hours. Timelapse imaging of spheroids to assess detachment was performed using a Leica Thunder Imager with a Okolab incubation chamber. Spheroids formed in round bottom ULA culture in the presence of conditioned media were transferred to a flat bottom ULA plate, treated with 0.5 mM calcein AM, and Z-stacked images were acquired every 15 minutes for 12 hours.

### Control trypsin assay

Cells were seeded onto 60mm dishes and allowed to adhere overnight before culturing in conditioned media for 72 hours. Adherent cells were subjected to controlled trypsinization (0.25% Trypsin/2.21 mM EDTA in HBSS without sodium bicarbonate, calcium & magnesium, 90 sec) and both adherent and detached cells were counted using a hemocytometer.

### CRISPR Drop out Screen

Human CRISPR Metabolic Gene Knockout library was a gift from David Sabatini (Addgene #110066; http://n2t.net/addgene:110066; RRID:Addgene_110066). Ovacr8 cells were infected with pooled CRISPR KO library to an MOI <0.3 to achieve ∼350-fold library coverage after selection with 1 µg/mL puromycin. The cells were passaged as needed every 2–3 days for 5, maintaining the full population. Cells were seeded and cultured in conditioned media for 72 hours before the controlled trypsin assay was performed as described above. The detached cell and adherent cell portions were collected separately in each condition and DNA was isolated using the Quick-DNA Miniprep Plus kit (Zymo Research, catalog no. D4069) following the manufacturer’s instructions. DNA from each condition pooled and used for gRNA amplification. To ensure adequate gRNA rep-resentation, calculations from Wohlhieter et al. ^134^ were used to achieve >250-fold coverage. The gRNA region was amplified using primers listed in **Table S7** and Ex Taq DNA polymerase (Takara, catalog no. RR001A) following the manufacturer’s protocol. The PCR conditions were as follows: one cycle at 94 °C for 2 minutes, followed by 30 cycles of 94 °C for 20 seconds, 63 °C for 30 seconds, and 72 °C for 45 seconds, concluding with a final extension step at 72 °C for 1 minute. The reaction was performed in a thermal cycler (Bio-Rad, T100™ Thermal Cycler, serial no. 621BR59583). The resulting PCR amplicons were purified using the Wizard® SV Gel and PCR Clean-Up System (Promega, catalog no. A9282) following the manufacturer’s instructions and pooled. The cleaned amplicons were sequenced using Next Seq v2.5 High Output 75 cycle on an Illumina NextSeq sequencer (see **Table S7** for primer details). Data was analyzed using the MAGeCK bioinformatics pipeline to identify differentially enriched genes^135^.

### Bioenergetic analysis of oxygen consumption rate (OCR) and extracellular acidification rate (ECAR)

The Agilent Seahorse XFp Metabolic Analyzer (Agilent, model S7802A) was used to assess mitochondrial respiration as described previously for adherent cells ^136^. Briefly, prior to the start of the experiment, cells were evenly seeded and cultured overnight in a Seahorse XFp cell culture plate (Agilent, 103022-100) at a density of 10,000 and 8000 cells/well for OVCA433 and SKOV3, respectively. The XFp sensor cartridge was hydrated in Agilent Seahorse XF Calibrant (Agilent,103022-100) at 37 °C in a humidified incubator (non-CO_2_) incubator overnight. On the day of the experiment, cell culture media was replaced with pre-warmed seahorse XF base RPMI media, pH 7.4 (Agilent,103576-100) supplemented with 1 mM sodium pyruvate (Agilent 103578-100), 2 mM glutamine (Agilent 103579-100), and 10 mM glucose (Agilent, 103577-100). Cells were then placed into a non-CO_2_ humidified incubator at 37 °C for 60 min. Mitochondrial stress test reagents (pharmacological manipulators of mitochondrial respiratory chain proteins) were diluted in pre-warmed XF assay media to achieve the following final concentrations in the cell culture well: 1.5 μM Oligomycin A (Sigma, 75351); 1 or 0.5 μM FCCP (Sigma, C2920) for OVCA433 and SKOV3, respectively; and 0.5 μM Antimycin A/Rotenone (Sigma, A8674,45656). Three basal rate measurements (3 min measurement time each) were taken prior to the injection of mitochondrial stress test reagents and three measurements of OCR/ECAR were obtained respectively following injection of compounds. Post-run, the cells were stained with crystal violet dye (0.05%) (Sigma-Aldrich, 229288) for seeding normalization. The dye was released from cells using 30% acetic acid and absorbance was measured at 590 nm, using GloMax Explorer (Promega) microplate reader.

### Immunofluorescence imaging

For immunofluorescence of adherent cells, cells were seeded at an equal density on coverslips or in Mattek dishes coated with poly-L-ornithine (Sigma, cat# P4957) and allowed to adhere overnight. Cells were stained live with CellMask deep red (ThermoFisher Scientific, cat# C10046) and fixed with 4% paraformaldehyde. Cells were washed four times with 1× PBS and either stained with 0.2 mg/mL filipin (Cayman Chemicals, cat# 70440) or permeabilized with 0.2% Triton X-100 in PBS for 5 min and then postfixed with 1% paraformaldehyde and 0.01% Tween 20 for 30 min. Permeabilized cells were blocked for 5 min with 3% BSA/PBS followed by incubation of corresponding primary antibody in 3% BSA/PBS for 1 h at room temperature. Prior to incubation with secondary antibody in 3% BSA/PBS for 1 h at room temperature, cells were washed three times with 1% Triton X-100 in PBS. Cells were then incubated with 0.15 µg/ml DAPI in 1× PBS for 1 min, washed three times with 1× PBS, mounted with fluorescence mounting medium (9 ml of glycerol [BP229-1; Fisher Scientific], 1 ml of 1× PBS, and 10 mg of p-phenylenediamine [PX0730; EMD Chemicals]; pH was adjusted to 8.0–9.0 using carbonate-bicarbonate buffer [0.2 M anhydrous sodium carbonate, 0.2 M sodium bicarbonate]) and sealed. 20X and 40X images were obtained at room temperature using a Nikon ECLIPSE Ti2 microscope with a 20×/0.75 N.A. or 40 ×/0.95 N.A. dry objective (Nikon DIC N2 Plan Apo) equipped with a camera (ORCA-Fusion C14440). Images were acquired using NIS-Elements AR software and processed using ImageJ. TIRF images were obtained at room temperature using a Nikon Ti-E with a Apo TIRF 100x 1.49 N.A. oil objective equipped with a camera (Photometrics Prime 95B). Images were acquired and processed using NIS-Elements software. Individual cells were segmented in the CellMask channel using Cellpose-SAM ^137^ followed by manual curation. Mean filipin intensity was quantified within segmented regions. Background was defined as the median intensity outside segmented cells and subtracted from the mean filipin signal.

### Chromatin immunoprecipitation

ChIP was performed as previously described in^130,138^ using the ChIP-grade antibodies described above. Cells were fixed in 1% paraformaldehyde for 5 min at room temperature and quenched with 1 mL of 2.5 M glycine for 5 min at room temperature. Cells were washed twice with cold PBS. Cells were lysed in 1 ml ChIP lysis buffer (50 mM Hepes-KOH, pH 7.5, 140 mM NaCl, 1 mM EDTA, pH 8.0, 1% Triton X-100, and 0.1% deoxycholate [DOC] with 0.1 mM PMSF and EDTA-free Protease Inhibitor Cocktail [P8340; Sigma-Aldrich]). Samples were incubated on ice for 10 min and then centrifuged at 3,000 rpm for 3 min at 4°C. The pellet was resuspended in 500 µl lysis buffer 2 (10 mM Tris, pH 8.0, 200 mM NaCl, 1 mM EDTA, and 0.5 mM EGTA with 0.1 mM PMSF and EDTA-free Protease Inhibitor Cocktail) and incubated at room temperature for 10 min. Samples were centrifuged at 3,000 rpm for 5 min at 4°C. Next, the pellet was resuspended in 300 µl lysis buffer 3 (10 mM Tris, pH 8.0, 100 mM NaCl, 1 mM EDTA, 0.5 mM EGTA, 0.1% DOC, and 0.5% N-lauroylsarcosine with 0.1 mM PMSF and EDTA-free Protease Inhibitor Cocktail). Cells were sonicated while on ice (10 s on, 50 s off). Next, 30 µl of 10% Triton X-100 was added to each tube, and samples were centrifuged at maximum speed for 15 min at 4°C. The supernatant was transferred to new tubes, and the DNA concentration was quantified. Samples were precleared for 1 h at 4°C on a rotator using 15 µl protein G Dynabeads (Thermo Fisher Scientific) in ChIP lysis buffer. Samples were centrifuged at maximum speed for 15 min at 4°C, after which the supernatant was transferred to a new tube. 50 µl of the antibody bead conjugate solution was added, and chromatin was immunoprecipitated overnight on a rotator at 4°C. The following washes were performed twice for 15 min each by rotating for 15 min at 4°C: ChIP lysis buffer, ChIP lysis buffer plus 0.65 M NaCl, wash buffer (10 mM Tris-HCl, pH 8.0, 250 mM LiCl, 0.5% NP-30, 0.5% DOC, and 1 mM EDTA, pH 8.0), and 10 mM Tris-HCl (pH 8.0) and 1 mM EDTA (pH 8.0). DNA was eluted by incubating the beads with fresh 50 mM Tris-HCl (pH 8.0), 10 mM EDTA (pH 8.0), and 1% SDS for 30 min at 65°C. Reversal of cross-linking was performed by incubating samples overnight at 65°C. Proteins were digested using 1 mg/ml proteinase K and incubating at 37°C for 5 h. Finally, the DNA was purified using the Wizard SV Gel and PCR Clean Up kit (A9282; Promega).

Immunoprecipitated DNA was analyzed by qPCR using PowerUp^TM^ SYBR^TM^ Green Mas-ter Mix (Applied Biosystems, cat#A25742). The following amplification conditions were used: 5 min at 95°C and 40 cycles of 95°C for 10 s and 30 s with 62°C annealing temperature. The assay ended with the following melting curve program: 15 s at 95°C, 1 min at 60°C, then ramping to 95°C while continuously monitoring fluorescence. All samples were assessed in triplicate. ChIP-qPCR primer sets are described in **Table S7**.

### Gene Set Enrichment Analysis from Shender VO, et al. cohort

Transcriptomic profiles of ovarian cancer cells incubated with autologous ascitic fluids from patients before and after chemotherapy was downloaded from Gene Expression Omnibus (GEO, GSE241909). Gene Set Enrichment Analysis (GSEA) was performed on the differential expression gene list using Metascape with the Wikipathways gene set database.

### Metabolite Measurement

Metabolites were measured by liquid chromatography-high resolution mass spectrometry adapted from previously published approaches ^139^. Conditioned media samples were sterile filtered through a 0.2 µm filter and snap frozen at −0 °C. Cell samples were quenched in 1 mL pre-chilled −0°C 80/20 methanol:water (v/v) and spiked with 50µL 1µM isotope labeled TCA cycle mix (Cambridge Isotope Laboratories MSK-TCA-A) pre-diluted in 80/20 methanol:water and 50µL 0.02 ng/µL Propionyl-L-carnitine-(N-methyl-d3) (Sigma Aldrich 52941). After vortexing for 1 min samples were returned to −0°C for 30 min, centrifuged at 18,000 x g 10 min at 4°C, and the supernatant was transferred to a deep well 96-well plate and evaporated to dryness under nitrogen gas. Samples were reconstituted in 100 µL and 2 µL of the sample was injected from a 4 °C autosampler into a ZIC-pHILIC 150 × 2.1 mm 5 µm particle size column (EMD Millipore) with a ZIC-pHILIC 20 x 2.1 guard column in a Vanquish Duo UHPLC System (Thermo Fisher Scientific) at 25 °C. Chromatography conditions were as follows: buffer A was acetonitrile; buffer B was 20 mM ammonium carbonate, 0.1% (v/v) ammonium hydroxide in water without pH adjustment, with a gradient of 0.5 min at 20% A then a linear gradient from 20% to 80% B; 20–20.5 min: from 80% to 20% B; 20.5–28 min: hold at 20% B at a 0.150 mL/min flow rate. Column elute was introduced to a Q Exactive Plus with a HESI II probe operating in polarity switching mode with full scans from 70-1000 m/z with an insource fragmentation energy of 1. Instruments were controlled via XCalibur 4.1 and data was analyzed on Tracefinder 5.1 using a 5ppm window from the predominant ion positive or negative. Area under the curves for each analyte was normalized to the matched internal standard or the nearest surrogate internal standard.

### Patient serum samples and clinical data

Pre-treatment (pre-surgery and pre-chemotherapy) blood was collected from patients with newly diagnosed epithelial ovarian cancer under University of Pittsburgh approved IRB protocols 20050019, 19070449, and 19060197. Blood was collected in red top tubes, centrifuged at 500g for 10 minutes, and serum was aliquoted into cryopreservation tubes and stored at −80°C until shipped to Metabolon for metabolomics analysis. Samples were shipped in a single batch to Metabolon to assess metabolite levels using the DiscoveryHD4 platform, which accurately identifies and quantitates 1,000+ metabolites with less than 5% process variability using 100uL of serum^140^. Samples were characterized using three independent platforms: ultrahigh-performance liquid chromatography tandem mass spectrometry (UPLC-MS/MS) in the negative ion mode, UPLC-MS/MS in the positive ion mode, and gas chromatography-mass spectrometry (GC-MS) after si-alylation. Compounds were identified based on chromatographic properties and mass spectra by comparing to a metabolomic library of purified standards. Clinical data were extracted from the electronic medical records by staff trained in medical record abstraction/coding (LB) and verified by comparison to the UPMC/HCC cancer registry database, which is accredited by the American College of Surgeons Committee on Cancer (ACoS-CoC). All patient primary diagnosis slides were reviewed by a single pathologist (EE) to confirm primary epithelial ovarian cancer diagnosis. Only patients completing a full course of primary platinum-based chemotherapy were included in this study.

### Quantification and Statistical Analysis

GraphPad Prism (version 10.0) was used to perform statistical analysis. Point estimates with standard deviations or standard errors were reported, as indicated, and the appropriate statistical test was performed using all observed experimental data. All statistical tests performed were two-sided and p-values < 0.05 were considered statistically significant. When necessary, outliers were identified and excluded by the ROUT method (Q = 0.1%).

## Notes

### Competing Interest Statement

The authors have declared no competing interest.

### Summary of Updates

Figures 1-5 revised; Figures S1-S5 revised; authors updated; supplemental files updates

